# Dietary stress induced macrophage metabolic reprogramming, a determinant of animal growth

**DOI:** 10.1101/2024.04.18.590077

**Authors:** Anusree Mahanta, Sajad Ahmad Najar, Nivedita Hariharan, Manisha Goyal, Ramaswamy Subramanian, Angela Giangrande, Dasaradhi Palakodeti, Tina Mukherjee

## Abstract

Nutrient sensing and signaling play pivotal roles in animal growth. However, under dietary stress, this system falters, leading to growth defects. While immune cells are increasingly recognized as key nutrient sensors, their impact on animal growth remains poorly understood. In this study, we investigate how *Drosophila* larval macrophages respond to excessive dietary sugar and identify a reconfiguration of their metabolic state. They undergo a glycolytic shift, intensify TCA activity, and elevate TAG synthesis. While typical of sugar-induced nutrient stress, these changes interestingly exert contrasting effects on animal growth: glycolysis and increased TCA activity inhibit growth, while the lipogenic shift promotes it. However, the lipogenic response is insufficient to counteract the metabolic events suppressing growth, resulting in an overall reduction in adult fly size under high sugar conditions. Stimulating a pro-lipogenic immune state facilitates growth recovery, suggesting a growth paradigm governed by immune-metabolic transitions. This study unveils the unexpected influence of macrophage metabolic reprogramming on organismal growth homeostasis during *Drosophila* development, highlighting immune cell states as central determinants of growth, particularly under dietary stress.

## Introduction

Body growth is a tightly regulated process that ensures formation of adults with correct size and proportions to finally influence survival and reproduction (Baron et al., 2015; Boulan et al., 2015; Nijhout et al., 2014). A complex integration of environmental and developmental cues governs the rate and duration of juvenile growth which determines the final adult body size (Penzo-Méndez and Stanger, 2015). It is also essential that these growth mechanisms are plastic to allow adaptation of developing animals to environmental challenges like infection and fluctuations in nutrition. The evolutionary conservation of mammalian growth control pathways in fruit flies has facilitated numerous studies revealing intricate communication between organs for systemic growth regulation in homeostasis and under varying environmental conditions (Koyama et al., 2020). Cross-organ communication among *Drosophila* nutrient sensor and responder tissues—including the fat body, brain, imaginal discs, muscle, and gut—is pivotal in regulating organ and body growth. Hormones, cytokines, and morphogens serve as the signaling molecules orchestrating this crosstalk (Reviewed in Chatterjee and Perrimon, 2021; Droujinine and Perrimon, 2016; Boulan et al., 2021). Understanding how these tissues sense environmental cues and adjust growth accordingly provides insights into the systemic growth control axis.

In this context, the functioning of the immune system with consequences on systemic growth is documented where examples of immune modulation and its impact on animal sizes have been described. Heightened immunity correlates with stunted growth while the opposite is true with animals with weak immune system (van der Most et al., 2011). The importance of maintaining immune cell numbers to enable systemic growth has also been recently described (Bakopoulos et al., 2020; Cho et al., 2020; Ramond et al., 2020). These evidences have alluded to immune cell functioning and its trade off with growth homeostasis. However, when it comes to growth axis, immune cells are seldom mentioned. Perhaps because these examples of growth modulation are described in conditions of infection, we consider changes in animal growth as consequence of altered immunity as opposed to their direct contribution to the larger scheme of developmental control of growth. The fact that immune cells are emerging as key nutrient sensors (Martínez-Micaelo et al., 2016; Newsholme, 2021) much like the fat body and brain, and implicated in developmental decisions (Juarez-Carreño and Geissmann, 2023), their role in growth homeostasis from a developmental standpoint does not seem unrealistic.

It is now increasingly appreciated that macrophages in response to their surrounding environment undergo metabolic-rewiring which in turn, determines their functional responses (Batista-Gonzalez et al., 2019; El Kasmi and Stenmark, 2015). The recent advances in high throughput transcriptomics and metabolic analysis have aided a deeper understanding of macrophage heterogeneity revealing distinct phenotypes that rely on metabolic pathways involving lactate (Geeraerts et al., 2021), purine (Li et al., 2022) and arginine (Viola et al., 2019). This is in addition to the already established M1 and M2 macrophage types employing aerobic glycolysis and fatty acid oxidation respectively (Galván-Peña and O’Neill, 2014). Whether these macrophage functional types are different subsets or one subset with potential for plasticity remains to be understood (Remmerie and Scott, 2018). Nonetheless, the link between the metabolic heterogeneity of macrophages and their functions has been widely implicated in both health and disease. Recent studies have in fact also shown *Drosophila* macrophage-like plasmatocytes to be highly heterogeneous with regard to adopting comparable metabolic remodeling (Cattenoz et al., 2020; Cho et al., 2020; Coates et al., 2021; Girard et al., 2021).

To that end, our work from the recent past has implicated *Drosophila* larval immune cells as regulators of animal growth (P et al., 2020). *Drosophila* blood cells, akin to myeloid cells (Evans et al., 2003), contributed significantly towards coordinating growth in conditions of dietary sugar stress. Growth retarding effects of excessive dietary sugar (high sugar diet, HSD) is evident from flies to mammals and the foremost underlying reason implicated in this pathological outcome is development of insulin resistance or inhibition of growth hormone signaling. We however found that animals with depleted number of immune cells grew poorly in conditions on dietary sugar stress. Intriguingly, animals with more active immune cells developed unexpectedly well on HSD and comparable to flies on regular diet. These findings highlighted immune cells as key modifiers of growth homeostasis in stress conditions. The work proposed immune cell state changes as a key paradigm for growth adaptation in stress conditions (P et al., 2020). Thus, immune control of growth both in homeostasis and in stress conditions which remains a poorly understood area, led us to take on board the current investigation. The immune underpinnings of systemic growth homeostasis, specifically with respect to growth retardation evident in high sugar intake forms the central focus of our investigation.

The present study employs a multi-prong, unbiased approach to explore immune-specific internal state changes that is central to growth control and adaptation on dietary stress conditions. Using a sensitized HSD model, the present study explores the immune-specific regulators that function to control animal growth in this form of excessive sugar linked over nutrition. We undertook a holistic methodology to ascertain immune-specific regulators that possibly promote or inhibit growth. The central and unexpected finding of this study is the impact of diet induced immune metabolic rewiring on organismal growth homeostasis. The findings from the work lead us to propose a key deterministic role played by immune cells in growth control and places the cellular immune system in the nexus of growth control paradigm.

## Results

### Dietary sugar overload impacts immune cell physiology and function

The central question of the study is to discern intracellular immune cell states governing body size control in a high sugar diet (HSD) induced stress. Therefore, we first characterized the status of immune cells themselves namely; cell numbers, basal metabolic state, morphology and function when exposed to HSD. To do this, we utilized the differentiating immune cell-specific marker Hemolectin, *Hml*^△^*>GFP* (*Hml*^△^*>GFP* crossed to *w^1118^)* background animals and exposed them to two different dietary regimes: one, short term exposure to HSD for four hours (referred to as 4hr.HSD, henceforth) as the means to gauge immediate changes induced in immune cells by short term intake of high sugar and second, a long-term, constitutive HSD feeding (referred as Ct.HSD, henceforth) to identify cell states established as a consequence of sustained high sugar intake by the animal. Specifically, for the short-term 4hr.HSD regime, *Hml*^△^*>GFP/w^1118^*feeding third instar larvae (72hr. AEL) rearing on regular food (RF, containing 5% sucrose) were transferred to HSD where they were allowed to feed for a brief period of four hours only. Subsequent to this, the larvae were dissected, bled and assessed for the aforementioned immune parameters. For Ct.HSD regime, *Hml*^△^*>GFP/w^1118^*embryos were collected and transferred to HSD, where the animals were reared until feeding 3^rd^ instar stage following which they were processed similar to 4hr.HSD, for immune characterization.

To assess HSD induced changes in immune cell numbers, we specifically monitored Hemolectin-positive (Hml^+^) and Hemolectin-negative (Hml^-^) cell populations across circulatory and sessile pools (see methods for further details on their assessment). For metabolic changes, we characterized immune cells for their intracellular redox state, glucose levels and lipid metabolism. For functional characterization, phagocytic bead uptake ability was measured (Hao et al., 2018) and finally for morphological changes, phalloidin stainings were undertaken to assess changes in cell morphology, size, shape, and length of immune cell filopodia extensions.

We observed that high sugar treatment severely impacted larval immune cell numbers in the long-term, Ct.HSD condition (Fig. 1a-d). We observed that while short-term, 4hr.HSD treatment to larvae did not reveal any changes in immune cell numbers, Ct.HSD animals, showed a significant decline in total immune cell numbers (Fig. 1c, c’). Specifically, a significant decline in sessile Hml^+^ population was apparent (Fig. S1 a-a’’’), while circulating cell numbers remained comparable across RF, 4hr.HSD and Ct.HSD (Fig. S1 b-b’’’). The number of Hml^-^ cells however remained comparable between RF, 4hr.HSD and Ct.HSD condition (Fig. 1d, Fig. S1a”’-b”’), which implied a specific sensitivity of the Hml^+^ cell population to high sugar exposure.

**Figure 1.**
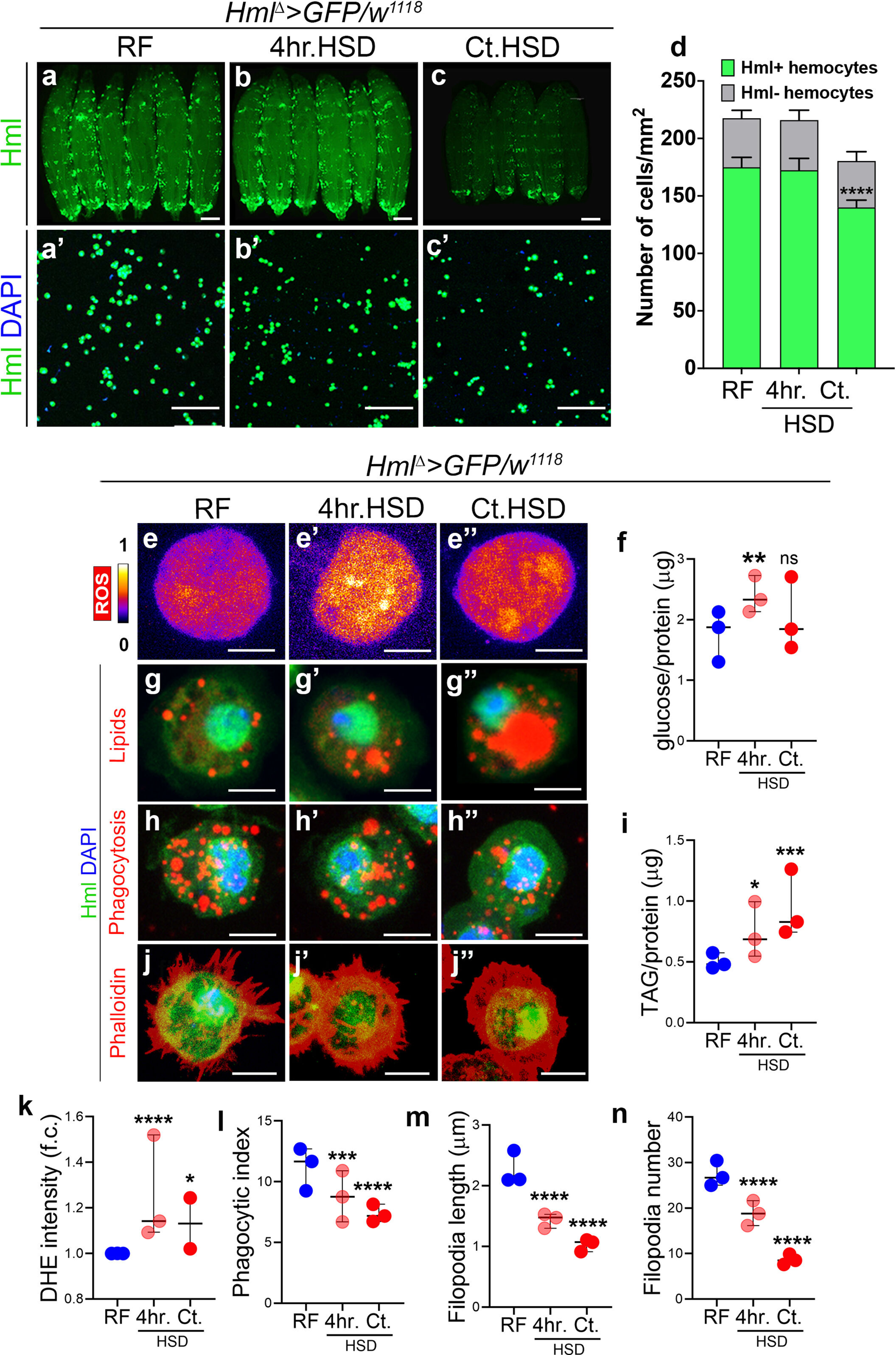
Dietary sugar stress affects larval macrophage physiology. Data information: DNA is stained with DAPI in blue, immune cells are marked in green (*Hml^Δ^>UAS-GFP*). DHE staining to assess ROS, is shown in spectral mode in panels (ee’’). Immune metabolic characterizations using nile red to mark lipids, bead uptake assay to assess phagocytosis and phalloidin to mark actin cytoskeletal changes is shown in red in panels (g-j”). Panels (a-c), scale bar is 1mm, (a’-c’), scale bar is 100μm and (e-j’’), scale bar is 5μm. The quantification in d represents mean with standard deviation. In quantification graphs (f, i, k, l, m and n), each dot represents individual experiments, extending from min. to max. Comparisons for significance are with regular food conditions and asterisks mark statistically significant differences (*p<0.05; **p<0.01; ***p<0.001; ****p<0.0001). The statistical analysis applied is Unpaired t-test with Welch’s correction or Two-way ANOVA with Dunnett’s multiple comparison test wherever applicable. RF, 4hr.HSD and Ct.HSD indicate conditions of larvae fed on regular food (RF), four hours high sugar diet (4hr.HSD) and constitutive high sugar diet (Ct.HSD) respectively. “N” is the total number of experimental repeats and “n” is the total number of larvae analyzed. See methods for further details on larval numbers and sample analysis for each of the experiments. (a-d) High sugar diet affects immune cell physiology. (a-c’) Representative images of feeding third instar larvae and immune cells on RF, 4hr.HSD and Ct.HSD. (a-a’) *Hml^Δ^>GFP/w^1118^*(*Control*, RF). (b-b’) *Hml^Δ^>GFP/w^1118^* (4hr.HSD) does not show any change in immune cell number as compared to *Control* (a-a’), (c-c’) *Hml^Δ^>GFP/w^1118^* (Ct.HSD) larvae show reduction in Hml-positive (Hml^+)^ immune cells when compared to *Control* (a-a’). (d) Quantification of total Hml^+^ immune cell numbers (green bars) in *Hml^Δ^>GFP/w^1118^* (*Control,* RF, N=3, n=18), *Hml^Δ^>GFP/w^1118^* (4hr.HSD, N=3, n=18, p=0.3790) and *Hml^Δ^>GFP/w^1118^* (Ct.HSD, N=3, n=18, p<0.0001). No change in Hml^-^ population is seen (grey bar). (e-e’’) Representative images of immune cells to assess ROS levels. Compared tp (e) ROS level in immune cell of *Hml^Δ^>GFP/w^1118^* (*Control*, RF), high sugar treated condition, (e’) *Hml^Δ^>GFP/w^1118^* (4hr.HSD) and (e’’) *Hml^Δ^>GFP/w^1118^* (Ct.HSD) show increased ROS levels. See quantification in k. Quantification of glucose levels in immune cells of *Hml^Δ^>GFP/w^1118^* (*Control*, RF, N=3, n=90), *Hml^Δ^>GFP/w^1118^* (4hr.HSD, N=3, n=90, p=0.0086) and *Hml^Δ^>GFP/w^1118^* (Ct.HSD, N=3, n=90, p=0.2911). Immune cells show higher glucose levels at 4hr.HSD as compared to *Control.* (g-g’’) Representative images of immune cells with Nile red staining to assess for lipid droplet accumulation. Compared to (g) *Hml^Δ^>GFP/w^1118^*(*Control,* RF), (g’) *Hml^Δ^>GFP/w^1118^* (4hr.HSD) and (g’’) *Hml^Δ^>GFP/w^1118^* (Ct.HSD) show gradual increase in immune cell lipid content. The immune cell in panel g’’ is a zoomed version of a cell selected from panel d’’ of Fig. S1. (h-h’’) Representative images of immune cells to assess phagocytosis through bead uptake assay. Compared to (h) *Hml^Δ^>GFP/w^1118^*(*Control*, RF), (h’) *Hml^Δ^>GFP/w^1118^* (4hr.HSD) and (h’’) *Hml^Δ^>GFP/w^1118^* (Ct.HSD) immune cells show reduction in number of internalised beads after 15 min of exposure. Quantifications in l. (i) Quantification of triglycerides (TAG) in immune cells of *Hml^Δ^>GFP/w^1118^* (*Control,* RF, N=3, n=105), *Hml^Δ^>GFP/w^1118^* (4hr.HSD, N=3, n=105, p=0.0209) and *Hml^Δ^>GFP/w^1118^* (Ct.HSD, N=3, n=105, p=0.0002). Immune cells show higher TAG levels on HSD both at 4hr. HSD and Ct.HSD, in comparison to *Control.* (j-j’’) Representative images of immune cells assessed for cellular morphology. Compared to (j) *Hml^Δ^>GFP/w^1118^* (*Control*, RF), (j’) *Hml^Δ^>GFP/w^1118^* (4hr.HSD) and (j’’) *Hml^Δ^>GFP/w^1118^* (Ct.HSD) show reduction in filopodia length (Quantifications in m) as well as filopodia number (Quantifications in n). (k) Quantification of ROS intensity levels of immune cells in *Hml^Δ^>GFP/w^1118^* (*Control,* RF, N=3, n=30), *Hml^Δ^>GFP/w^1118^*(4hr.HSD, N=3, n=30, p<0.0001) and *Hml^Δ^>GFP/w^1118^*(Ct.HSD, N=2, n=20, p=0.0232). (l) Quantification of bead uptake assay of immune cells in *Hml^Δ^>GFP/w^1118^* (*Control,* RF, N=3, n=30), *Hml^Δ^>GFP/w^1118^*(4hr.HSD, N=3, n=30, p=0.0001) and *Hml^Δ^>GFP/w^1118^*(Ct.HSD, N=3, n=30, p<0.0001). (m) Quantification of immune cell filopodia length in *Hml^Δ^>GFP/w^1118^*(*Control*, RF, N=3, n=30), *Hml^Δ^>GFP/w^1118^*(4hr.HSD, N=3, n=30, p<0.0001) and *Hml^Δ^>GFP/w^1118^*(Ct.HSD, N=3, n=30, p<0.0001). (n) Quantification of immune cell filopodia number in *Hml^Δ^>GFP/w^1118^* (*Control,* RF, N=3, n=30), *Hml^Δ^>GFP/w^1118^*(4hr.HSD, N=3, n=30, p<0.0001) and *Hml^Δ^>GFP/w^1118^*(Ct.HSD, N=3, n=30, p<0.0001).

Next, we compared the metabolic states and assessed for ROS levels by dihydroethidium (DHE) staining and observed that it was elevated with high sugar exposure in 4hr.HSD and Ct.HSD immune cells (Fig. 1 e-e’’, k and Fig. S1 c-c’’). Biochemical means to estimate glucose levels revealed a significant rise in immune cell glucose following the short term 4hr.HSD exposure, which however was not detected in the long-term Ct.HSD regime (Fig 1 f). This implied that immune cell glucose levels increased immediate to sugar exposure, but gradually plateaued in the longer term.

We observed that high sugar treatment also led to an overall increase in lipogenesis, much more evident in the longer-term Ct.HSD than 4hr.HSD condition (Fig. 1g-g”, i and Fig. S1 d-d”, g-g”). For lipid measurements, we employed nile red staining to mark lipid droplets, TAG biochemical measurements to assess total TAG and *UAS-LSD2-GFP* genetic reporter line to assess their lipogenic state (Fauny et al., 2005). Specifically, we observed a gradual increase in the number of lipid droplets (Fig. 1 g-g’’ and Fig. S1 d-d’’) and total TAG level in immune cells from 4hr.HSD and Ct.HSD when compared to RF condition (Fig. 1 i). These signatures of increasing levels of TAG in the cells corroborated with their lipogenic potential as seen with increasing *LSD2-GFP* reporter expression (Fig. S1 g-g’’). The data highlighted the sensitivity of immune cells to HSD, and the overall impact on their internal lipid homeostasis when faced with dietary sugar stress. Altogether, the increased glucose levels following sugar exposure and the gradual increase in lipogenesis were suggestive of induction of metabolic programs to accommodate excessive sugar (Musselman et al., 2013).

Functionally, excessive sugar exposure severely impaired immune cell phagocytic abilities. A gradual decrease in number of internalised beads was evident in the immune cells. This was seen as early as in 4hr.HSD treatment and dramatically reduced in Ct.HSD condition (Fig. 1 h-h’’, l and Fig. S1 e-e’’). Morphologically, compared to numerous filopodia seen protruding from the immune cell surface from RF larvae (Fig. 1 j and Fig. S1 f), a reduction in the number and length of filopodia was evident even at 4hr.HSD exposure which was further pronounced in immune cells when subjected to Ct. HSD condition (Fig. 1 j-j’’, m, n and Fig. S1 f-f’’).

The overall temporal profiling of immune cell number, cytoskeleton dynamics, phagocytosis and metabolism revealed manifestation in metabolic and functional capabilities in immune cells as early as 4hr of HSD feeding. A clear decline in immune cell phagocytic ability with increased lipogenesis was evident and these changes exaggerated with longer exposure to dietary sugar. Any relevance of such sugar induced immune cell state changes on adult body size, was investigated next. To do this we undertook an unbiased genomewide RNA*i* screening approach as the means to identify specific candidates whose functioning in immune cells affected animal growth on HSD.

### Identification of immune-specific growth modulators using an *in-vivo* RNA*i* genetic screening approach

Setting up of the unbiased whole genome RNA*i* screen was conducted with a total of 1052 RNA*i* strains which were specifically expressed in Hml^+^ differentiating immune cells of the *Drosophila* larvae using the *Hml^Δ^-Gal4* (*Hml^Δ^>*) as the driver line (Sinenko and Mathey-Prevot., 2004). Specifically, males from each RNA*i* lines were crossed to virgin *Hml^Δ^>UAS-GFP* females on regular food and from the crosses, 35-40 embryos were collected, transferred to high sugar diet and reared at 29°C until adult flies eclosed whose sizes were thereafter scored. In each experimental set, the RNA*i* crosses were tested in two batches as biological replicates for enhanced accuracy. For comparison to quantify changes in adult sizes, *Hml^Δ^>UAS-GFP*/*w^1118^*adults grown on RF and on HSD conditions were used as controls that demonstrated normal adult fly size and growth retarded HSD flies respectively (Fig. 2a).

**Figure 2.**
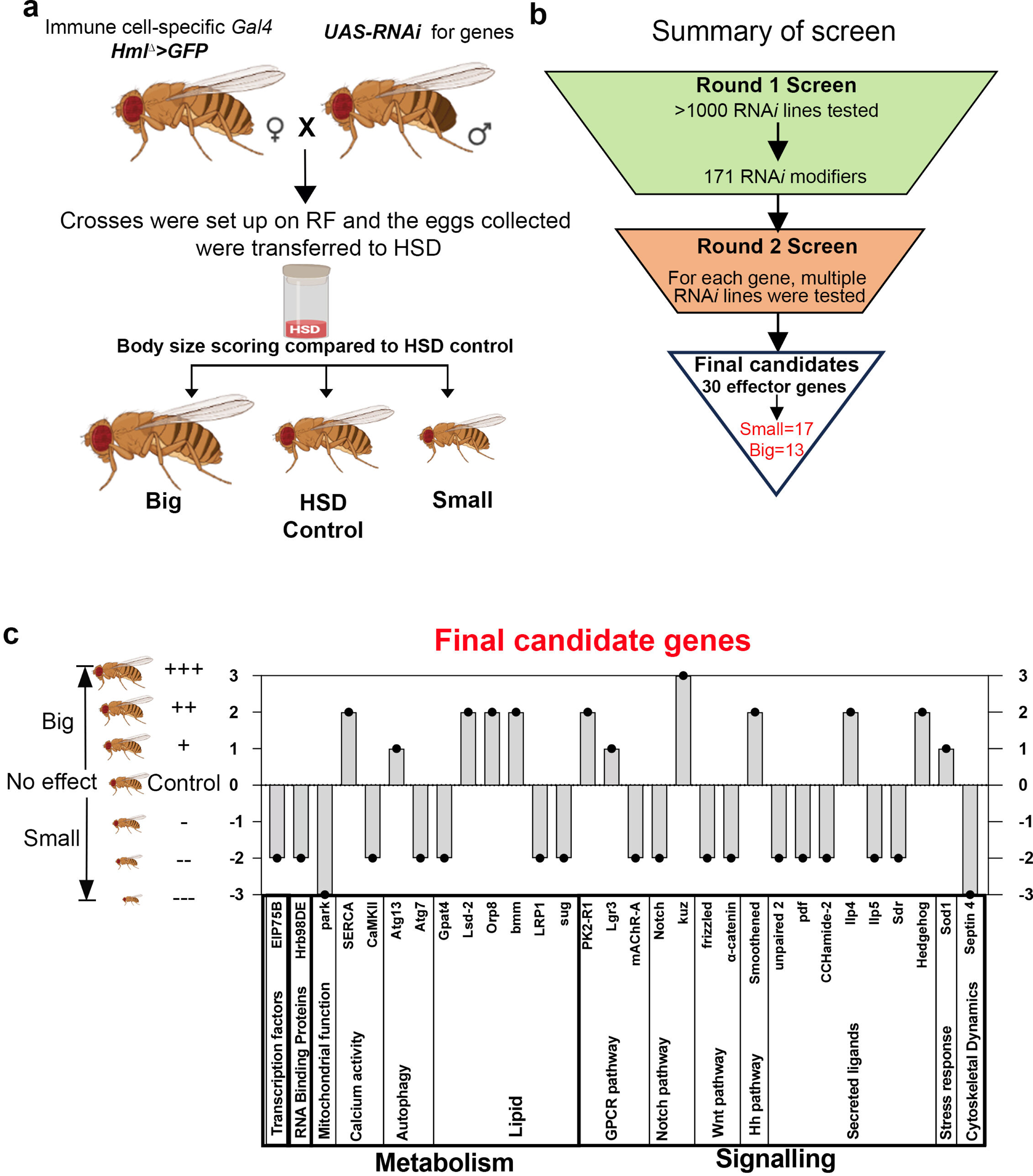
Identification of immune-specific modulators of animal growth in dietary sugar stress condition. (a) Schematic representation of *in-vivo* genome wide RNA*i* screen. Females of immune cell specific Gal4 driver (*Hml^Δ^>GFP*) line were crossed to *UAS-RNAi* males. Eggs collect-ed on regular food were transferred to high sugar diet (HSD). Eclosed flies were scored for body size phenotype. (b) Summary of the results from *in-vivo* RNA*i* screen. In the first round, >1000RNA*i* lines were tested. Modifiers obtained from first round were further tested in second round with multiple RNA*i* lines (See Table S1 for details on the lines tested). Finally, we arrived at 30 effector genes with 17 genes being positive regulators of growth and 13 as negative regulators. (c) Summary of the final effector/candidate genes obtained from the screen. The genes were categorized based on their biological functions. “Big” and “Small” phenotypes, were graded into mild (+) or (-), moderate (++) or (--) or severe (+++) or (---) categories.

For scoring, one day old adult flies obtained post eclosion were phenotypically screened and scored for their body size. To increase robustness of the screening process, the size scoring was performed independently by three different individuals in a blind unbiased manner. As an initial body size scoring paradigm, the adults from the RNA*i* crosses were scored either for any further reduction in their size than seen in HSD flies and were marked “Small”, or any recovery in their size if they appeared any closer to size seen in regular food raised controls and were marked “Big” respectively (Fig. 2a).

Based on this assessment criteria, in first round of RNA*i* screen, a total of 171 RNA*i* lines were identified as “modifiers” of adult size on HSD condition. Of these, 101 lines showed size reduction and were smaller than HSD control adults, implying that these lines are positive regulators of growth on HSD condition. Very interestingly, 18 RNA*i* strains restored the adult size defect seen in HSD. The emerging adults from these RNA*i* crosses, were larger in size than observed in HSD controls and were rather closer in their size towards RF wild type controls. These “Big” genes were designated as negative regulators of growth. The remaining 797 RNA*i* lines did not demonstrate any deviation from HSD controls and were recorded as “no effect” (NE) and were listed as non-modifiers (Fig. 2b).

For the total 171 modifier lines identified from Round 1, we next undertook a second round of screening. Here, we tested multiple RNA*i* lines against each candidate and only those candidates where we observed consistent growth phenotypes across two or more RNA*i* lines were finally selected. Majority of the RNA*i* lines chosen here are published lines validated for their function (Table S6). Depending on the extent of size modulation observed they were further graded. For “Big” and “Small” phenotypes, they were also graded into mild (+) or (-), moderate (++) or (--) or severe (+++) or (---) categories (Fig. 2c and Table S1).

We found a total of 30 genes that showed consistent phenotype with more than one RNA*i* line and were listed as “final candidate genes” (Fig. 2c and Table S1). Of these 17 were identified as “Small” and 13 were identified as “Big” and majority of these lines were moderate modifiers of the growth phenotype while only a few were mild effectors. This finding implied a robust contribution by the immune cells on growth in HSD condition. Subsequently, Flybase was used to determine the known or predicted functions of these genes (Fig. 2c and Table S1). When functionally categorised, these top 30 candidate genes came under the categories that included diverse cellular functions ranging from transcription factors to metabolic and signaling genes (Fig. 2c).

As the major cohort important for animal growth, the signaling genes seemed expected, however, we observed an unexpected influence on growth in this category. Signaling pathway components of the Notch, Wnt and JAK/STAT pathway, were identified necessary for growth as their down regulation in immune cells caused a retardation in adult fly size compared to HSD controls. Hedgehog (Hh) signaling, contrarily was identified as a negative modulator of growth. Interestingly, both *hh* and and its receptor, *smoothened* (Alcedo et al., 1996) appeared in the screen and blocking their expression in blood cells, led to growth recovery. The adults were much larger in size than the corresponding HSD controls. The data implied that in HSD condition, immune cells signaling exerted dual control on growth, where players like Notch, Upd2, and Wnt enabled growth, but Hh and it’s signaling inhibited growth.

The other biological process that was over-represented, was the “metabolism category”. Under this, majority encoded functions related to “lipid metabolism”. Specifically, lipogenic genes like *Glycerol-3-phosphate acyltransferase 4* (*Gpat4)*, involved in triglyceride synthesis (Heier and Kühnlein, 2018) and transcription factors, *sugarbabe (sug),* O*xysterol receptor protein 8 (Orp8)* (Kokki et al., 2021; Mattila et al., 2015; Repa et al., 2000), known to promote lipogenic expression on high sugar diet were identified as positive growth regulators. Importantly, the screen also identified *Brummer, bmm*, a key lipolytic gene (Grönke et al., 2005), whose down regulation, showed growth recovery with adult flies much larger in size than HSD control. These data revealed a significant role for immune lipid levels on systemic growth and implied growth promoting functions for immune lipogenesis but growth inhibitory consequences for immune cell lipid turnover (Fig. 2c).

Overall, the findings from the screen, revealed unexpected opposing states in the Hml^+^ population of immune cells where they were growth enabling with players like *Notch, upd2, Gpat4* but also disabling with respect to *hh, smo* and *bmm*. Apart from these, other candidates also identified included members of, mitochondrial metabolism (*parkin*), autophagy (*Atg13* and *Atg7*) and cytoskeletal remodeling (*Septin4*) all of which were identified as positive regulators. These altogether highlighted an important contribution of immune intrinsic state in systemic coordination of animal growth.

At this point, it also seemed necessary to ascertain any association of changes in immune cell numbers with body size phenotype. When assessed for a few randomly selected candidate genes from the screen, we found that knocking down these genes and the corresponding change on immune cell numbers did not correlate with their adult fly size phenotype (Fig. S2 a-e). For instance, loss of *Sod1* in immune cells, while it caused a reduction in immune numbers as evident by fewer *Hml^Δ^>UAS-GFP* positive cells, this genetic condition was identified as “Big” in the screen (Fig. S2b, and Table S1). Similar for *Lsd2^RNAi^*, whose expression in immune cells, increased their numbers, was also identified as “Big” in the screen (Fig. S2c and Table S1). On the contrary, loss of *Septin1* or *park* in immune cells even though they did not show any dramatic difference in cell numbers (Fig. S2d, e), the adults in these conditions were identified as Small” phenotype (Fig. 2c and Table S1). Moreover, the modulation of these genes in the immune cells did not alter larval growth, as the mutant larva were comparable to HSD control larvae in terms of their overall sizes (Fig. S2 a-e). Only when assessed for adult fly sizes, they were affected (Tables S1). These evidences further reinstated and confirmed our previously reported finding that immune cell state rather than numbers determined growth and that immune cells specifically contributed towards adult growth regulation and not larval growth homeostasis (P et al., 2020).

### Transcriptional changes induced by HSD highlights reprogramming of macrophage metabolic state

The physiological changes (Fig.1) together with the distinct impact on growth seen upon modulating lipogenic and lipolytic genes as identified in the screen (Fig. 2), highlighted a relevant contribution of immune cell metabolic state on systemic growth. This led us to dwell deeper and gain a holistic understanding of metabolic state changes invoked in immune cells in response to HSD. For this, we undertook a genome wide transcriptomics analysis by RNA sequencing of immune cells (Fig. S3a). As a side-by-side comparison, whole larval transcriptome was also undertaken to comprehend global changes invoked by HSD as opposed to specific changes initiated only in the immune cells (Fig. S3a). Total immune cells from 4hr.HSD and Ct.HSD fed larvae were collected and processed for bulk RNA sequencing with RF immune cells as control sample (Fig. S3a). Transcriptomics analysis of whole larvae was also performed in these conditions (Fig. S3a).

The biological processes influenced by 4hr.HSD, and longer-term Ct.HSD, using Gene Ontology (GO) analysis highlighted immediate transcriptional changes that remained persistent even in constitutive HSD condition (Tables S2, 3). These included down regulation of genes encoding JAK-STAT signaling pathway, Toll/Imd, ecdysone signaling and Wnt signaling pathway. Along with these, genes involved in cell migration, cell-matrix adhesion and integrin signaling were also seen down regulated upon HSD treatment. The transcriptional changes converged with morphological analysis shown in Fig.1 and implied that high sugar transcriptionally dampened their immune potential and cytoskeletal remodeling proteins. We also assessed the expression of some of the screen candidates, like *Upd2* (JAK/STAT Pathway), *Fz*, *α-catenin* (Wnt-signaling pathway), but did not observe any transcriptional alteration. Nevertheless, the overall transcriptional down regulation of the aforementioned signaling pathways highlighted their sensitivity to excessive sugar and the associated implication on growth identified in the screen, corroborated with their functional requirement.

When assessed for up regulated pathways, metabolic processes emerged as the most over-represented/predominant biological process (Fig. 3 and Table S2). Specifically, lipid metabolism was highlighted here as well, where up regulation of lipogenic genes was observed (Fig. 3a). This includes *Acetyl CoA carboxylase (ACC)*, the rate limiting enzyme involved in *de novo* fatty acid synthesis (Parvy et al., 2012), *GPAT, AGPAT* and and *Lipin*, (Heier and Kühnlein, 2018) all enzymatic members of the TAG synthesis pathway (Fig. 3 and Table S2, 3). *ACC* up regulation was only seen in 4hr.HSD condition (Fig. 3b) but not with long term chronic exposure in Ct.HSD (Tables S2 and S3). Genes like *Gpdh1* and *Lipin* were up regulated at 4hr.HSD condition and continued to be over-expressed even in longer term HSD exposure (Fig 3b and Table S2, 3). A unique GO term the “fatty acid biosynthetic process” which included genes predicted in fatty acid elongase activity (*CG8534, CG9459, CG30008* and *CG331100*) were also seen up-regulated but only in the longer term Ct.HSD condition (Table S3), which implied changes in carbon length of the fatty acids in larval immune cells with constitutive sugar exposure. Immune cells also showed a significant increase in the expression of beta-oxidation pathway enzyme *Acyl-CoA synthetase (Acsl)*, but compared to lipogenic genes, the lipid breakdown candidates were not over-represented in the transcriptional landscape. This revealed a transcriptional reprogramming of the immune cells towards a lipogenic state.

**Figure 3.**
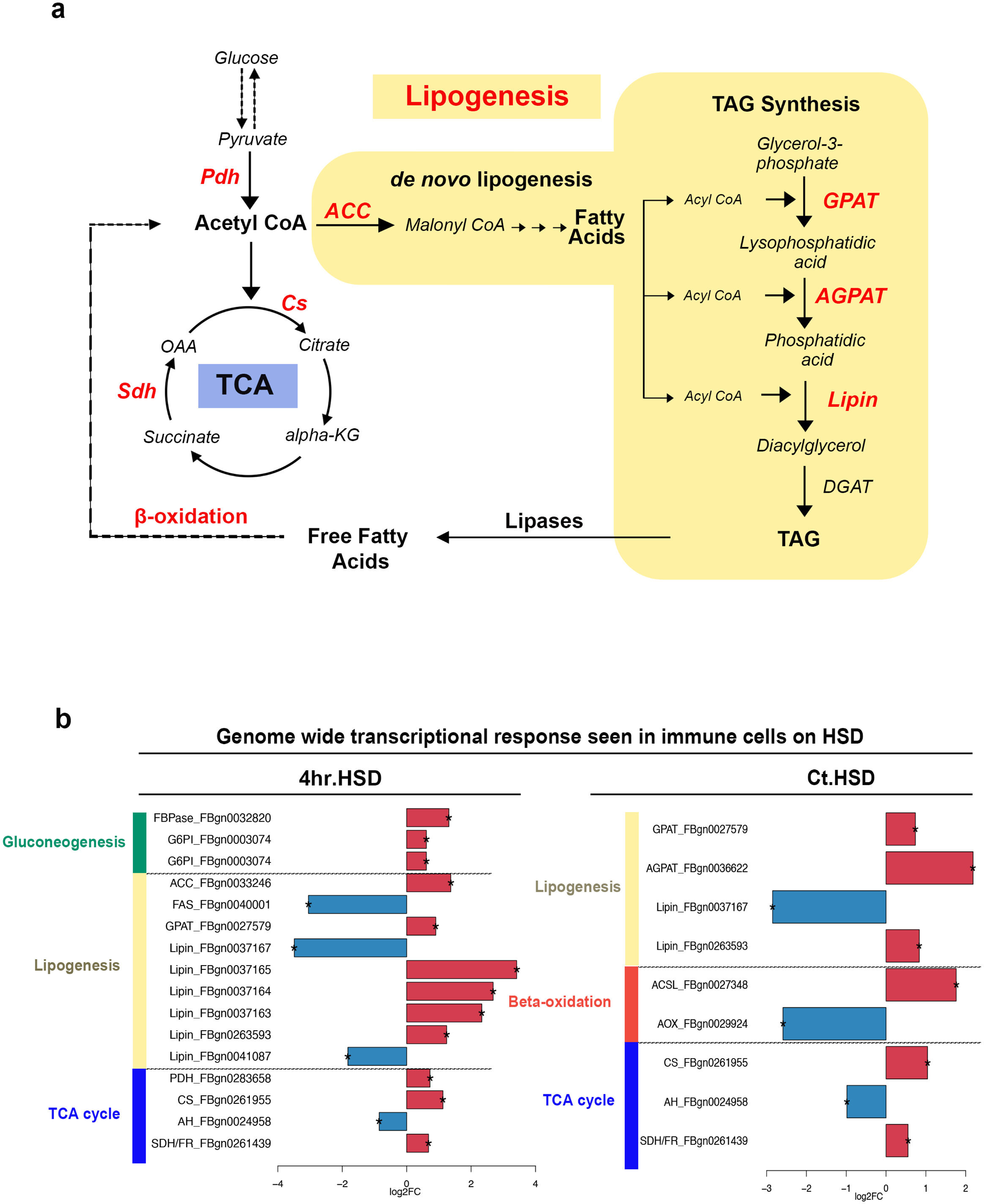
Dietary sugar stress induces metabolic rewiring in immune cells. (a) Diagrammatic representation of overall transcriptional changes seen in metabolic genes in immune cells on HSD with short term (4hr.HSD) and long-term (Ct.HSD) exposure. All genes shown in red indicate their transcriptional up regulation, this includes TCA enzymes, *de novo* lipogenesis, TAG synthesis pathway and beta-oxidation enzymes. (b) Bar plots show temporal changes in respective metabolic genes, and their paralogs. Red bars indicate up regulated genes, and blue bars indicate down-regulated genes. *ACC,* which is a key de novo lipogenic enzyme, is seen up regulated immediately upon HSD exposure, but not with long-term exposure, when only TAG synthesis is seen up regulated. beta-oxidation enzyme, *acyl-CoA synthetase long-chain (ACSL)* is however seen upregulated in constitutive HSD condition, but not in 4h, short term.

In addition to lipid, genes/ enzymes of pyruvate metabolism and cycling like *Malic enzyme b (Men-b)* and *Mitochondrial pyruvate carrier (Mpc1)* were also seen up regulated, indicative of increased pyruvate metabolism, which was expected given the high sugar impact (Table S2). Corroborating with the involvement of pyruvate, the TCA pathway components were also seen to be up regulated (Fig. 3 a, b). This was interpreted based on genes like *midline uncoordinated (muc)* with *Pyruvate dehydrogenase (Pdh)-*like activity*, Citrate synthase 1 (Cs1)* and *Succinate dehydrogenase* (*Sdh).* They were all up regulated in high sugar conditions (Fig. 3 a, b). Mitochondrial pyruvate dehydrogenase moderates TCA activity by regulating pyruvate entry into the TCA cycle (Leiter et al., 1978; Linn et al., 1969), and with *muc* which has PDH like activity (Marygold, S.J, 2024), it’s up regulation at 4hr alluded to heightened pyruvate entry into the TCA (Fig. 3). Its expression was however not up-regulated in the long term Ct.HSD exposure (Fig. 3b), but the sustained activation of other TCA genes, *Cs* and *Sdh* (Fig. 3 a, b) implied heightened TCA-activity.

We also observed up-regulation of genes necessary for “glutathione metabolism” (Tables S2, 3). This corroborated with TCA activity increase and induction of ROS in HSD immune cells shown in Fig. 1e-e’’. A small subset of genes involved in “regulation of cell division” which included genes for spindle assembly and cytokinesis and “actin cytoskeleton reorganization” were also seen up regulated (Table S2, 3) and reinstated the impact of sugar stress on cellular proliferation and functions (Fig. 1d).

Overall, the transcriptomic analysis furthered our understanding of the metabolic events induced in immune cells upon HSD exposure. The data revealed metabolic reprogramming in immune cells upon sensing of high sugar. These include induction of heightened TCA activity as early as 4hr post excessive sugar exposure and a shift towards a lipogenic state (Fig. 3 a, b). The early identification of *ACC*, and sustained up regulation of TAG synthesis genes, *GPAT1*, *AGPAT* and *Lipin* corroborated with the increased lipogenesis seen in the aforementioned physiological characterizations (Fig. 1f-f’). These metabolic changes were very specific to the blood tissue and not evident in the animal as a whole (Fig S4). In the animal on the other hand, an overall dampening of metabolism was observed (Fig. S4a) which was more clearly seen with long-term HSD exposure (Fig. S4b). Developmental genes were rather seen to be up regulated in the whole animal on HSD (Tables S4 and S5) thereby implying that immune cells rewired their metabolism distinctly to enable processes to deal with incoming sugar load and invoked specific internal events not globally initiated.

The outcomes from the genetic screen allude to contribution of immune metabolic states on systemic growth homeostasis. Specifically, the identification of lipogenic and lipolytic genes as moderators of growth from the screen led us to investigate the additional metabolic changes identified in the transcriptomic data and their contribution to growth in dietary sugar stress. To conduct this, we employed metabolic and genetic approaches and dissected the specifics of each step in a systematic manner.

### Heightened TCA activity and glycolytic shift in HSD immune cells: Metabolic states that repress growth

As the first step, we addressed TCA and employed liquid chromatography-tandem mass spectrometry (LC-MS/MS) to confirm the observed transcriptomics changes in TCA activity. This was followed by genetic approaches to modulate corresponding TCA genes to assess the impact of animal growth on HSD condition. In this stage of our analysis, we conducted a rather quantitative approach to score body sizes. We chose wingspan areas and fly body length estimations (Lee et al., 2008, 2004) as a proxy for estimating the extent of changes brought about by corresponding genetic manipulations on animal growth.

We performed metabolic flux analysis with isotopic U^13^C-pyruvate (Buescher et al., 2015., Jang et al., 2018) to discern any changes in the rate of TCA activity. To achieve this, immune cells from regular food (RF) and constitutive high sugar diet (Ct.HSD) conditions were incubated with U^13^C-pyruvate, and the flow of C13 into TCA cycle intermediates was assessed. Pyruvate enters the TCA cycle via pyruvate dehydrogenase (PDH), where it is converted into acetyl-CoA, and this contributes to two carbons into the TCA metabolites (Fig. 4a). Pyruvate incorporates three carbons in oxaloacetate (OAA) via pyruvate carboxylase (PC), which further adds on to citrate and thus contributes to the TCA cycle. Pyruvate is also converted into lactate via LDH and contributes to all the three carbons of lactate. Thus, the differential labelling of carbons in TCA metabolites and lactate was considered as a measure of change in pyruvate flux under RF and Ct.HSD condition. Apart from metabolic flux analysis, we also conducted steady-state targeted comparative analysis of TCA cycle metabolites. For this, immune cells were isolated from animals raised in RF and Ct.HSD exposure and processed for steady-state metabolite analysis.

**Figure 4.**
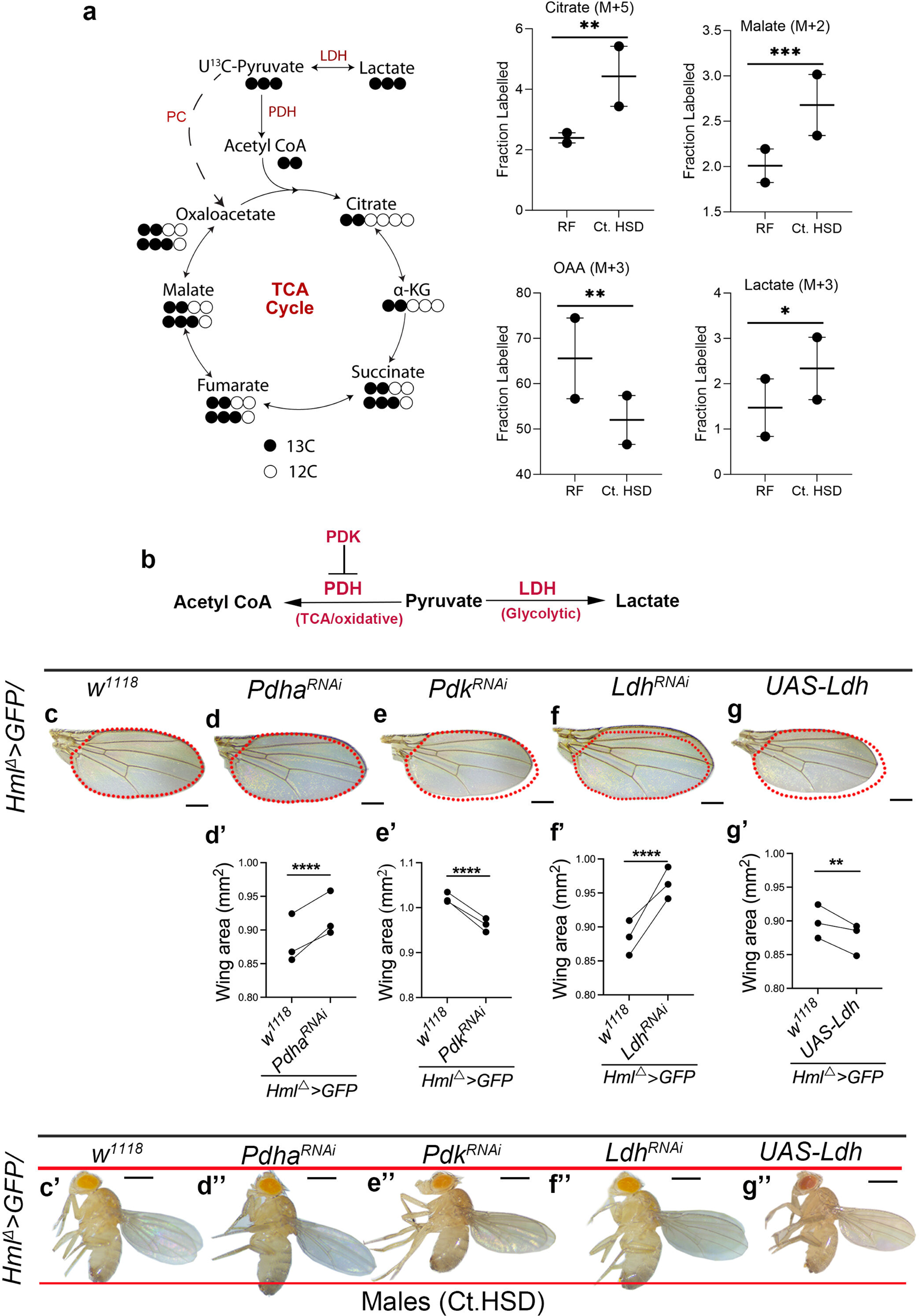
An oxidative and aerobic glycolytic state in immune cells represses growth on HSD. Data information: Scale bar: 0.5mm for flies and 0.25mm for wings. In quantification graphs, shown in panel (a), (d’-g’) each dot represents an experimental repeat. Except for panel (a), where comparions are with respect to Control on RF, in all other panels comparison for significance is with respect to Control on HSD. Asterisks mark statistically significant differences (*p<0.05; **p<0.01; ***p<0.001; ****p<0.0001). The statistical analysis applied is Two-way ANOVA, main effects test. “N” is the total number of repeats. “n” is the total number of animals (larvae or adult flies) analyzed. Only right wing from each adult fly was selected for quantification. The red dotted line in the panels marks the wing span area and is used as a reference to showcase any change in wing span area across genotypes. The two horizontal red lines in the panels is used as a reference to showcase any change in fly lengths across genotypes. RF and Ct.HSD correspond to regular food and constitutive high sugar diet respectively. (a) Distribution of labeled U^13^C pyruvate in TCA metabolites and lactate in *Hml^Δ^GFP>/w^1118^* (Control, RF) and *Hml^Δ^GFP>/w^1118^* (Ct.HSD) conditions. Ct.HSD led to an increase in M+5 label incorporation in citrate (N=2, n=55, p=0.0018), M+2 label incorporation in malate (N=2, n=60, p=0.0007) and a decrease in M+3 label incorporation in OAA (N=2, n=55, p=0.0028). Increase in M+3 label incorporation in lactate was also noticed (N=2, n=70, p=0.0337). (b) Pyruvate metabolism into acetyl CoA under the regulation of PDH enzyme fuels the TCA /oxidative metabolism. PDK inhibits PDH activity and regulates TCA. Pyruvate conversion to lactate is driven by LDH enzymatic activity. (c-g’’) Modulating larval immune cell TCA and glycolytic activity affects adult growth. Representative images of fly wings of adult males (c-g) showing size phenotype on Ct.HSD from respective genetic backgrounds. Compared to (c) Ct.HSD *Control* (*Hml^Δ^>GFP/w^1118^*), moderating TCA activity by (d) expressing *Pdha^RNAi^* (*Hml^Δ^>GFP/Pdha^RNAi^*) to reduce TCA led to increase in animal size while (e) *Pdk^RNAi^* (*Hml^Δ^>GFP/Pdk^RNAi^*), to increase TCA activity, decreased their size furthermore. (f) Down-regulating immune glycolytic activity by expressing *Ldh^RNAi^* (*Hml^Δ^>GFP/Ldh^RNAi^*) lead to size increase while (g) over-expressing *Ldh* (*Hml^Δ^>GFP/UAS-Ldh*) led to decrease in animal sizes. (d’-g’) Wing span quantifications in males. (d’) *Hml^Δ^>GFP/Pdha^RNAi^* (N=3, n=95, p<0.0001 in comparison to corresponding Ct.HSD *Control*, *Hml^Δ^>GFP/w^1118^*, Ct.HSD, N=3, n=76), (e’) *Hml^Δ^>GFP/Pdk^RNAi^* (N=3, n=77, p<0.0001 in comparison to corresponding Ct.HSD *Control, Hml^Δ^>GFP/w^1118^,* Ct.HSD, N=3, n=66), (f’) *Hml^Δ^>GFP/Ldh^RNAi^* (N=3, n=125, p<0.0001 in comparison to Ct.HSD *Control*, *Hml^Δ^>GFP/w^1118^,* Ct.HSD, N=3, n=101) and (g’) *Hml^Δ^>GFP/UAS-Ldh* (N=3, n=122, p<0.0064 in comparison to Ct.HSD *Control, Hml^Δ^>GFP/w^1118^,* Ct.HSD, N=3, n=70). (d’’-g’’) Representative images of adult males on Ct.HSD from respective genetic backgrounds compared to control (c’).

The levels of TCA cycle metabolites between RF and Ct.HSD failed to show any difference in the steady state conditions (Fig. S5a). The isotopic metabolite measurements however revealed increased flux of pyruvate into TCA metabolites and lactate under HSD conditions (Fig. 4a and Fig. S5b). Specifically, our isotopic labelling data showed increased, higher 13C label incorporation in citrate upon HSD, which indicates the increased pyruvate flux towards TCA cycle (Fig. 4a and Fig. S 5b). Moreover, malate also showed an increase in M+2 label incorporation in the HSD condition which is donated by PDH mediated entry of pyruvate into the TCA (Fig. 4a and Fig. S 5b). These data showed that immune cells change their metabolic state from less oxidative to more oxidative upon high sugar exposure. The rate of PC metabolism in HSD condition was however reduced as decrease in M+3 labelling in OAA was seen. This could be attributed to the concomitant rise in pyruvate entry into TCA via PDH (Fig. 4a and Fig. S5b). An increased flow of labelled C13 pyruvate into lactate in HSD condition was also apparent (Fig. 4a and Fig. S 5b). Even though *Ldh* transcript levels did not reveal any significant up regulation, the increased M+3 labelling in lactate upon HSD exposure implied increased LDH activity and demonstrated an aerobic glycolytic shift in these immune cells (Vander et al., 2009). Thus, considering the biochemical data, increased sugar exposure in the immune cells, exaggerated the overall flow of pyruvate into the TCA via PDH and into lactate via LDH (Fig. 4a).

Next, we modulated TCA cycle and LDH activity to comprehend the precise contribution of these metabolic state changes on growth regulation on HSD, respectively. To gauge TCA function, we genetically modulated PDH by expressing RNA*i* against the *Pyruvate dehydrogenase E1 alpha subunit (Pdha)* and down regulated its expression in immune cells (*Hml^△^>GFP/Pdha^RNAi^*) and assessed animal growth in HSD condition. This genetic manipulation showed a mild but significant recovery in animal size as apparent in their wingspan areas (Fig. 4c-d’’). When assessed for body lengths although males were larger compared to HSD controls (Fig. 4c’-d’), the overall significance was indifferent and may be due to underlying inconsistencies in batches (Fig. S6a). The quantitative analysis however did not reveal any difference in the females (Fig. S6 f-f’’’). The converse experiment, to further increase TCA activity in immune cells, by down regulating pyruvate dehydrogenase kinase *(Pdk)* enzyme, the negative regulator of PDH (Bowker-Kinley et al., 1998) however led to dramatic reduction in animal sizes (Fig. 4e-e’’ and Fig. S6 b and g-g’”). Here, a consistent reduction in body length was evident across males (Fig. S6b) and females (Fig. S6g’’’). Altogether, the data implied a growth inhibitory role for heightened immune cell TCA-activity, but given the mild recovery seen on blocking *Pdha* and its inconsistency across males and female genders, suggested additional inputs that operated towards growth impairment beyond the TCA cycle.

We modulated pyruvate conversion towards lactate and performed similar genetic manipulations of *Ldh* enzyme and assessed for adult growth. We observed that knockdown of *Ldh* enzyme increased adult fly sizes both in terms of wing span (Fig. 4f-f’ and Fig. S6 h-h’) and body lengths (Fig. S6 c and h’’’) in males and females. Converse experiments with over expression of *Ldh* in immune cells, caused a further reduction in animal size in comparison to HSD-control adults (Fig. 4g-g’’ and Fig. S6 d, i-i’’’). These data unlike PDH genetic manipulations unveiled a stronger influence of immune cell glycolytic shift on animal size control. The recovery in adult fly sizes seen with down regulating *Ldh* in immune cell, indicated the sufficiency of immune cell lactate production on adult growth inhibition in HSD.

### Opposing effects of immune cell *de novo* lipogenic and lipolytic state in systemic growth regulation

The bulk transcriptome revealed up regulation of *de novo* and TAG lipogenic enzymes and the screen revealed a growth enabling role for immune cell *GPAT* function and a growth inhibitory for *bmm*. We therefore undertook real-time quantitative PCR method to confirm the expression levels of all these lipogenic pathway enzymes and *bmm*. As done for *RNA*-seq, immune cells from larvae reared on 4hr.HSD, Ct.HSD and RF control were isolated and assessed. Here as well, a significant up regulation of *ACC*, was detected in 4hr.HSD immune cells but not with constitutive treatment (Fig. S7a1). *GPAT1, 1-Acylglycerol-3-phosphate O-acyltransferase 4 (Agpat4), Lpin* and *Midway* encoding *acyl coenzyme A: diacylglycerol acyltransferase (DGAT1)* was analyzed and we found that *GPAT1* and *Midway* were up regulated in the long-term Ct.HSD condition (Fig. S7a2-a5). Additionally, expression of *Glycerol 3 phosphate dehydrogenase1 (Gpdh1),* that *catalyzes* the conversion of dihydroxyacetone phosphate to glycerol-3-phosphate and provides the backbone for TAG synthesis was also seen up-regulated in Ct.HSD condition (Fig. S7a6). Expression analysis of *bmm*, failed to show any difference in its transcript levels (Fig. S7 a7). These collectively confirmed an early transcriptional shift towards increasing intracellular immune cell lipid synthesis, through *de novo* fatty acid synthesis early and through TAG synthesis as the longer-term response to high dietary sugar exposure.

We genetically deconstructed their roles and first addressed the contribution of the *de novo* lipogenic pathway (Fig. 5a). We investigated the expression dynamics of ACC protein in immune cells on RF (Fig. 5b), 4hr.HSD (Fig. 5c) and Ct. HSD (Fig. 5d) condition. Unlike the RT-PCR results, no dramatic increase in ACC protein was seen in 4hr.HSD condition and was comparable to RF control (Fig. 5c compared to 5b, Fig. 5e). The ACC protein levels however declined in Ct.HSD condition and was much lower than seen in RF (Fig. 5d compared to 5b, Fig. 5e). This implied that the initial transcriptional induction of *de novo* lipogenic enzyme did not lead to a corresponding up-regulation in its protein levels. With sustained high sugar exposure, the levels of ACC declined furthermore. We know that HSD condition drives intracellular lipid levels to increase (Fig. 1g-g”). The impact of ACC on any changes in immune-cell lipid levels was therefore characterized. We observed that down regulation of *ACC* expression in larval immune cells (*Hml^△^>GFP/ACC^RNAi^*), led to a significant reduction in lipid levels than seen in HSD control immune cells (Fig. 5f-g, Fig. S7b-c). These data implied a role for ACC function in immune lipogenesis and suggested that even though levels of ACC in long term HSD conditions was not high, its activity contributed towards increasing lipid levels. Next, we assessed adult fly sizes, and observed that immune down regulation of *ACC* (*Hml^△^>GFP/ACC^RNAi^*) led to smaller animal sizes in comparison to HSD controls (Fig. 5k-l’’ and Fig. S7g, k-l’’’). This was evident in wingspan (Fig. 4l-l’ and Fig. S7 l-l’) and body length quantifications (Fig. S7 g and l’’’) across male and female adult flies. We also did the converse experiment and over expressed *ACC* enzyme in immune cells (*Hml^△^>GFP/UAS-ACC*) which led to a further increase in intracellular lipid levels, much more than seen in HSD control immune cells (Fig. 5h and Fig. S7d). Moreover, this genetic manipulation led to a significant increase in fly sizes across genders (Fig. 5m-m’’ and Fig. S7 h, m-m’’’), which suggested that even though ACC function was necessary to support growth it was limited in its capacity. We speculate the inability of immune cells to maintain *ACC* expression in long-term sugar exposure restricted the extent of its growth function. We speculate that the increased PDH activity in immune cells is opposed by ACC function that exerts an additional route to divert sugar derived acetyl CoA intermediate into lipids (Fig. 3a). While the sugar breakdown is growth inhibitory, the ACC mediated *de novo* lipogenic arm confers adaption to growth on HSD. The gain of ACC also suggests that favoring *de novo* lipogenesis enables better growth recovery on HSD.

**Figure 5.**
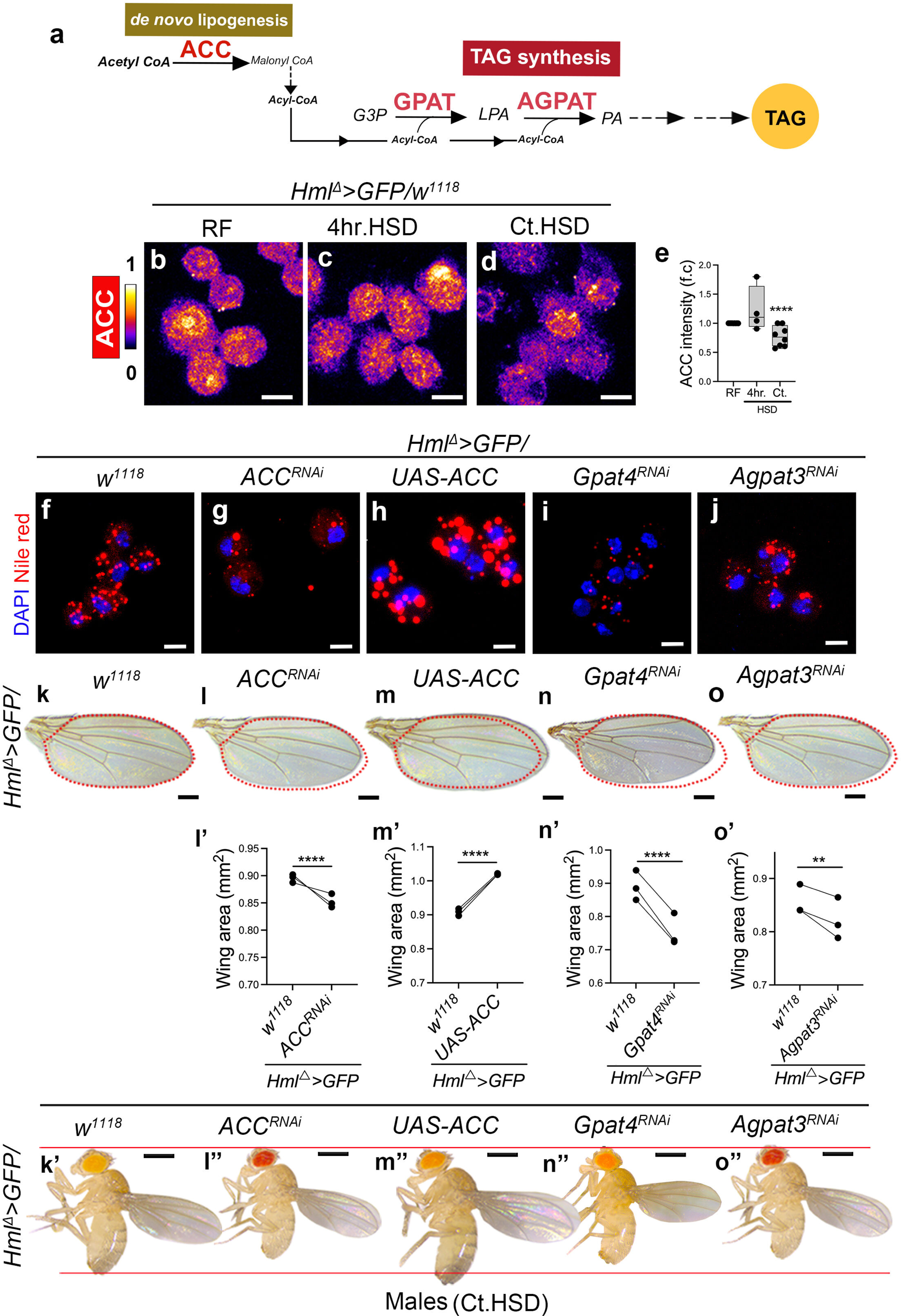
Immune cell lipid homeostasis and systemic growth regulation on HSD. Data information: DNA is stained with DAPI (blue). ACC staining is shown in spectral mode in panels (b-d). Nile red staining to mark lipids is shown in red in panels (f-j). Scale bar: 5μm for immune cells, 0.5mm for flies and 0.25mm for wings. In quantification graphs (e), (l’-o’) each dot represents an experimental repeat. Except for panel (e), where comparions are with respect to Control on RF, in all other panels comparison for significance is with respect to Control on HSD. Asterisks mark statistically significant differences (*p<0.05; **p<0.01; ***p<0.001; ****p<0.0001). The statistical analysis applied is Two-way ANOVA, main effects or with Dunnett’s multiple comparison test wherever applicable. “N” is the total number of repeats. “n” is the total number of animals (larvae or adult flies) analyzed. Only right wing from each adult fly was selected for quantification. The red dotted line in the panels marks the wing span area and is used as a reference to showcase any change in wing span area across genotypes. The two horizontal red lines in the panels is used as a reference to showcase any change in fly lengths across genotypes. RF and Ct.HSD correspond to regular food and constitutive high sugar diet respectively. See methods for further details on larval numbers and sample analysis for each of the experiments. (a) Schematic representation of *de novo* lipogenesis and Triacylglycerol (TAG) synthesis pathway. (b-d) Representative images of immune cells stained to visualize Acetyl CoA carboxylase (ACC) protein expression. Compared to ACC protein levels in (b) RF Control (*Hml^Δ^>GFP/w^1118^*), (c) 4hr.HSD *Control* (*Hml^Δ^>GFP/w^1118^*) showed no change in ACC expression, while (d) Ct.HSD *Control* (*Hml^Δ^>GFP/w^1118^*) the expression of ACC is decreased dramatically and is much reduced than levels seen in RF condition (b). (e) Relative quantification of ACC protein expression, RF (N=9, n=30), 4hr.HSD (N=4, n=30) and Ct.HSD (N=8, n=30, P<0.0001). (f-j) Representative images of immune cells stained to observe lipid droplets (Nile red, red) in (f) Ct.HSD *Control* (*Hml^Δ^>GFP/w^1118^*) and with immune-specific (g) loss of *ACC* function, (*Hml^Δ^>GFP/ACC^RNAi^*), (h) gain of *ACC* expression (*Hml^Δ^>GFP/UAS-ACC*), (i) loss of *Gpat4* (*Hml^Δ^>GFP/Gpat4^RNAi^*) and (j) loss of *Agpat3*(*Hml^Δ^>GFP/Agpat3^RNAi^)* function. Compared to Ct.HSD *Control* (f), loss of immune cells lipid synthesis both *de novo* (g) or TAG synthesis (i) and (j) led to reduced lipid droplets in them. Contrarily, gain of *de novo* lipid synthesis (h) shows increased lipid droplets in them. Please see Fig. S7 (b-f) for immune cells marked in green (*Hml^Δ^>UAS-GFP*) along with nile red. (k-o’’) Modulating larval immune cell lipid homeostasis affects adult growth. Representative images of fly wings of adult males (k-o) showing size phenotype on Ct.HSD from respective genetic backgrounds. Compared to (k) Ct.HSD *Control* (*Hml^Δ^>GFP/w^1118^*), (l) loss of *ACC* function (*Hml^Δ^>GFP/ACC^RNAi^*) leads to growth retardation, while (m) gain of immune *ACC* expression (*Hml^Δ^>GFP/UAS-ACC*) shows growth recovery as the flies are larger than HSD *Control* adults. Similarly, loss of TAG synthesis, by blocking (n) *Gpat4* (*Hml^Δ^>GFP/Gpat4^RNAi^*) or (o) *Agpat3* (*Hml^Δ^>GFP/Agpat3^RNAi^*) shows reduction in animal size. (l’-o’) Quantification of wingspan in (l’) *Hml^Δ^>GFP/ACC^RNAi^*(N=3, n=90, p<0.0001 in comparison to Ct.HSD *Control*, *Hml^Δ^>GFP/w^1118^*, N=3, n=71), (m’) *Hml^Δ^>GFP/UAS-ACC* (N=3, n=75, p<0.0001 in comparison to Ct.HSD *Control*, *Hml^Δ^>GFP/w^1118^*, N=3, n=102), (n’) *Hml^Δ^>GFP/Gpat4^RNAi^* (N=3, n=92, p<0.0001 in comparison to Ct.HSD *Control*, *Hml^Δ^>GFP/w^1118^*, N=3, n=79) and (o’) *Hml^Δ^>GFP/Agpat3^RNAi^* (N=3, n=56, p=0.0078 in comparison to Ct.HSD *Control*, *Hml^Δ^>GFP/w^1118^,* N=3, n=48). (l’’-o’’) Representative images of adult males on Ct.HSD from respective genetic backgrounds compared to control (k’).

Next, we did quantitative assessment of the TAG synthesis pathway. Down regulation of TAG lipogenic pathway components, *Gpat4* (*Hml^△^>GFP/Gpat4^RNAi^*) and *Agpat3* (*Hml^△^>GFP/Agpat3^RNAi^*) in immune cells led to decreased lipid levels in immune cells (Fig. 5i-j and Fig. S7e-f). Their down regulation confirmed the smaller animal size phenotype seen in the screen, but this was now clearly evident in wingspan areas (Fig. 5n-o’ and Fig. S7n-o’) and body length quantifications (Fig. 5n’’-o’’ and Fig. S7 i, j, n’’-o’’’). This was observed consistently across males and females and altogether reinstated the importance of TAG synthesis in immune cells to drive animal growth on HSD condition.

While the former process contributes to *de novo* TAG synthesis, we also know that immune cells express lipid scavenging receptors (Franc et al., 1996) which can add to intracellular lipid levels. In this context, *croquemort (Crq)*, the CD-36 homolog in flies (Franc et al., 1996, Guillou et al., 2016) its role in immune cell lipid uptake and high fat induced dietary stress outcomes is well described (Kiran et al., 2022). Protein levels of Croquemort demonstrated an increase in immune cells on Ct.HSD treatment (Fig. S8 a-c). Its genetic down regulation in immune cells led to reduction in their intracellular lipids (Fig. S8d-e’), and a retardation in adult fly size (Fig. S8 f-k) in both males and females. The data demonstrated that along with *de novo* lipogenesis, lipid uptake also contributed towards increasing intracellular TAG levels and these processes together enabled growth on HSD. Inhibiting either pathway was sufficient to impair the growth promoting benefits of the other lipogenic steps. The use of immune cells and their lipogenic potential in HSD condition was a prominent outcome from these analyses.

Post lipogenesis, lipolysis drives TAG breakdown (Fig. 6a) and the cycle of lipid synthesis coupled to breakdown maintains intracellular lipid homeostasis (Huang et al., 2014). The screen identified *bmm* dependent TAG breakdown, as a negative regulator of animal growth for which we conducted similar quantifications. Even though the RT-PCR expression analysis failed to show any difference (Fig. S7a7), genetic down regulation of *bmm* (*Hml^△^>GFP/bmm^RNAi^*), led to further elevation in intracellular immune cell lipid levels on HSD (Fig. 6 b-c and Fig. S9 a, b), which implied its active state in immune cells. Down regulation of *bmm,* was accompanied by an increase in animal size (Fig. 6 e-f” and Fig. S9 d-e’’). Contrarily, *bmm* over expression (*Hml^△^>GFP/UAS-bmm*) in the immune cells not only led to reduction in intracellular lipid levels (Fig. 6d, Fig. S9c), but also caused further retardation of animal sizes than seen in HSD controls (Fig. 6 g-g’’ and Fig. S9 f-f’’’). This implied a deleterious impact of immune-specific lipid breakdown on animal growth and alluded to immune-derived free fatty acids and their debilitating effects on systemic growth homeostasis in HSD.

**Figure 6.**
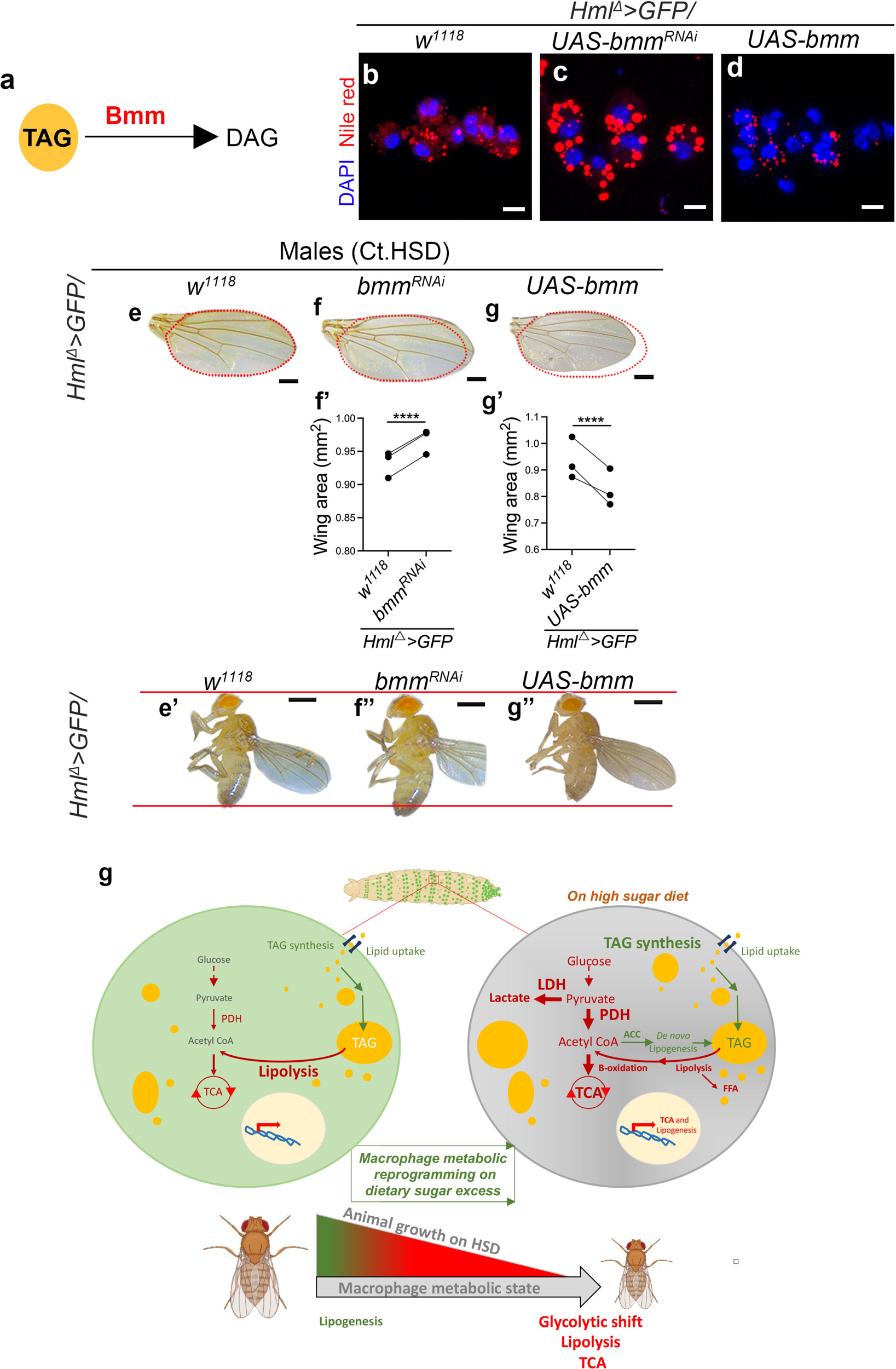
Immune cell lipolytic state as inhibitor of systemic growth on HSD. Data information: DNA is stained with DAPI (blue), immune cells are shown in green (*Hml^Δ^>UAS-GFP*). Scale bar: 5μm for immune cells, 0.5mm for flies and 0.25mm for wings. In quantification graphs (f’) and (g’), each dot represents an experimental repeat. Comparisons for significance are with Control on HSD and asterisks mark statistically significant differences (*p<0.05; **p<0.01; ***p<0.001; ****p<0.0001). The statistical analysis applied is Two-way ANOVA, main effects test. “N” is the total number of repeats. “n” is the total number of animals (larvae or adult flies) analyzed. Only right wing from each adult fly was selected for quantification. The red dotted line in the panels marks the wing span area and is used as a reference to showcase any change in wing span area across genotypes. The two horizontal red lines in the panels is used as a reference to showcase any change in fly lengths across genotypes. RF and Ct.HSD corresponds to regular food and constitutive high sugar diet respectively. (a) Bmm lipase enzyme function breaksdown Triacylglycerol (TAG) into diacylglycerol (DAG). (b-d) Representative images of immune cells on Ct.HSD stained to show lipid droplets (Nile Red, red). Compared to lipid levels seen in (b) Ct.HSD *Control* (*Hml^Δ^>GFP/w^1118^*), (c) loss of immune cell *brummer* function (*Hml^Δ^>GFP/bmm^RNAi^*) led to increase in lipid droplets and (d) gain of *bmm* expression (*Hml^Δ^>GFP/UAS-bmm*) led to decrease in lipids respectively. Please see Fig. S9 (a-c) for immune cells marked in green (*Hml^Δ^>UAS-GFP*) along with nile red. (e-g’’) Modulating larval immune cell lipolysis affects adult growth. Representative images of fly wings of adult males (e-g) showing size phenotype on Ct.HSD from respective genetic backgrounds. Compared to (e) Ct.HSD *Control* (*Hml^Δ^>GFP/w^1118^*), (f) loss of *bmm* (*Hml^Δ^>GFP/bmm^RNAi^*) led to recovery in adult fly size while (g) increase in *bmm* expression (*Hml^Δ^>GFP/UAS-bmm*) in immune cells caused a further reduction in size. (f’-g’) Quantification of wingspan in (f’) *Hml^Δ^>GFP/bmm^RNAi^* (N=3, n=96, p<0.0001 in comparison to Ct.HSD *Control*, *Hml^Δ^>GFP/w^1118,^* N=3, n=132) and (g’) *Hml^Δ^>GFP/UAS-bmm* (N=3, n=91, p<0.0001 in comparison to Ct.HSD *Control*, *Hml^Δ^>GFP/w^1118,^* N=3, n=56). (f’’-g’’) Representative images of adult males on Ct.HSD from respective genetic backgrounds compared to control (e’). (f) High sugar diet induced macrophage metabolic reprogramming in *Drosophila* larvae affects animal growth. Compared to regular diet, where immune cells are oxidative, internalise lipids and are lipolytic, HSD stress induces transcriptional rewiring of their metabolic state. They show elevated pyruvate entry into the TCA cycle and into lactate formation. They also undergo a lipogenic shift (green arrows, lipid droplets in yellow) which is mediated by *de novo* lipogenesis (ACC dependent), lipid uptake and elevated TAG synthesis. The resulting TAGs are broken down through the lipolytic pathway (Free Fatty Acids, FFA) and this overall maintains the levels of TAG in the HSD immune cells. The shift to glycolysis, increased TCA and lipolysis is growth inhibitory (steps marked in red arrows), but the lipogenic metabolic shift favours growth (steps marked in green). The extent of lipogenesis is however not sufficient to oppose the metabolic events that lead to growth repression, which leads to the overall reduction of adult fly size on HSD. The importance of lipids inside immune cells remains a key to cope up with dietary stress induced growth inhibition.

### Upstream modifier of immune cell lipogenesis

Overall, the data lead us to conclude that the exposure to high sugar diet induced increased pyruvate metabolism into driving TCA and a corresponding glycolytic shift. While these may be intrinsic to the immune compartment to counter high sugar stress, the outcome of these catabolic activities is a systemic inhibition of growth. The induction of TAG lipogenesis in these cells, through *de novo* and lipid uptake however functions to favor growth. Although our findings in the current state cannot discriminate between the strengths of different lipogenic modalities and their precise contribution in enabling growth, ACC whose functions allow diversion of sugars into lipids (Cao et al., 2008; Yore et al., 2014), its role in capacitating growth on high sugar appears central. Downstream of lipid synthesis, lipid breakdown is growth inhibitory and most likely adds to components like free fatty acids or acetyl-CoA which are known drivers of excessive sugar induced lipotoxicity (Postic and Girard, 2008; Unger, 2003). These results lead us to speculate that the lipogenic and lipolytic balance in immune cells is key to countering growth inhibitory stress imposed by HSD. As the balance shifts towards *de novo* lipogenesis, the growth achieved is superior than seen on HSD.

Regulation of *de novo* lipogenic enzyme ACC therefore appears critical. Given the limiting expression levels of *ACC* on HSD, it led us to explore upstream players that could be involved in this. For this we took to screen candidates and addressed a few of them for their effects on immune cell lipid levels. Of all the players, Notch and Hedgehog (Hh) components were identified as strong regulators of growth, where Notch signaling emerged as a positive regulator of growth while Hh signaling impaired growth on HSD. Thus, we undertook characterization of these two signaling pathways. We assessed for their activity and subsequent impact of their modulation on immune cell lipid levels and animal growth.

We found that Notch activity was dramatically down regulated in immune cells obtained from HSD treated animals (Fig. S10 a-c). Notch activity was assessed by measuring levels of intracellular domain of Notch protein (NICD), which was detected using an antibody against NICD. The reduction in immune NICD levels implied sensitivity of Notch signaling in these cells to high sugar intake. Contrary to this, expression analysis of Hh pathway in immune cells using Ci protein expression as a corresponding read out of Hh activity (Chen et al., 1999), revealed a significant increase in Ci expression in immune cells on HSD (Fig. S10 d-f). This implied differential impact of excessive sugar intake on immune signaling, where high dietary sugar down regulated Notch activity but led to stabilisation of Hh signaling in them.

We explored the impact of down regulating Notch signaling by expressing a dominantnegative version of the corresponding S3 cleavage enzyme, Presenilin (Psn, *Hml^△^>GFP/UAS-Psn^DN^*, Fortini, 2001) in the immune cells. Psn, catalyzes the intramembrane cleavage of Notch receptor and we found that expression of dominant negative *Psn*, led to reduction in animal size, both in terms of wing span and body lengths (Fig. S10 g-h’’ and Fig. S11 a, d-e’’’). Contrarily, expression of Notch-activated form in immune cells (*Hml^△^>GFP/UAS-N^Act^*), as the means to activate Notch signaling in them was sufficient to rescue the size defect seen in HSD condition (Fig. S10 i-i’’ and Fig. S11 b, f-f’’’). On the other end, consistent with genetic screen data, down regulation of *hh* in the immune cells (*Hml^△^>GFP/hh^RNAi^*) led to recovery in HSD adult fly sizes (Fig. S10 j-j’’ and Fig. S11 c, gg’’’). Unlike Notch, which consistently affected both genders, loss of hedgehog signaling showed stronger recovery in female sizes (Fig. S11 g-g’) than that seen in males (Fig. S10 j-j’). Although not conclusive the emerging differences in male and female sizes in few of the backgrounds like PDH, suggest sex-specific regulation of growth on HSD and the differential control exerted by immune cells on this axis.

Finally, we assessed for any change in intracellular lipid content and observed that immune cells with reduced Notch signaling in Ct.HSD, showed reduction in their lipid droplet content (Fig. S10 k-l’). Raising Notch activity on the other hand, (*Hml^△^>GFP/Notch^act^)* increased intracellular lipid levels (Fig. S10 m-m’). This suggested that intra-cellular Notch activity was critical towards driving a lipogenic state in immune cells. Intriguingly, a similar increase in immune cell intracellular lipid level was also observed upon loss of *hh* function in them (Fig. S10 n-n’). When assessed for any changes in ACC expression in immune cells, down regulating Notch although did not reveal any change in its levels (Fig. S10 op), increasing Notch signaling was sufficient to raise ACC protein levels (Fig. S10 q,s). Down regulating *hh* in immune cells was also sufficient to raise ACC protein levels in HSD immune cells (Fig. S10 r, s). Collectively, these data support a lipogenic role for Notch, and an anti-lipogenic role of Hh signaling in immune cells.

The data lead us to propose an immune state, where the differential impact of sugar on signaling pathways like Notch and Hh most likely limits the extent of their intracellular lipogenic potential. Thus, even though lipogenesis is induced, its insufficiency to counter growth inhibitory metabolic steps invoked by TCA, glycolytic shift and lipolysis is perhaps why the animal emerges as a smaller adult on high sugar diet.

## Discussion

### Dietary stress induced macrophage metabolic reprogramming, a determinant of animal growth trajectory

This research underscores the intricate link between the metabolic states of macrophages and their influence on animal growth. Through a comprehensive approach involving immune cell characterization, genome-wide RNA*i* screening, transcriptomics, morphometrics, and metabolomics, we elucidate the nuances of macrophage metabolic adaptations in response to excessive dietary sugar and their repercussions on animal size regulation (see Fig. 6g). Our findings reveal an unexpected utilization of immune cells’ lipogenic potential as the means to promote systemic growth under high sugar diet conditions.

Previous studies have established larval circulating immune cells, akin to macrophages, (Evans et al., 2003) as primarily lipid-scavenging entities, actively engaging in lipolysis and oxidative metabolism (Cattenoz et al., 2020). However, we observe a notable shift in their metabolic profile in the presence of dietary sugar excess. Larval immune cells undergo heightened tricarboxylic acid (TCA) cycle activity, induce a glycolytic switch, and ramp up triacylglycerol (TAG) synthesis. Importantly, the induction of the lipogenic state is intricately regulated by lipolytic processes, maintaining intracellular lipid levels. These metabolic adaptations likely confer immune cell tolerance to excessive sugar-induced stress but concurrently impacts systemic growth dynamics. Through genetic characterization, we identify steps in TCA cycle activity, glycolytic transition, and increased lipolysis as growthinhibitory, whereas the induction of lipogenesis and TAG synthesis promotes growth. Thus, in high sugar, an imbalance between growth favoring and growth-inhibiting pathways results in an anti-growth immune-metabolic state, leading to growth retardation under high sugar diet conditions (refer to Fig. 6g). Genetic interventions inhibiting either the growth-opposing immune metabolic pathways or promotion of growth-favoring metabolic processes suffices to restore growth levels comparable to those observed under a regular diet. This study unravels a novel regulatory mechanism wherein immune metabolic alterations exert precise control over systemic growth dynamics. Despite the positive effects of lipogenesis, the metabolic state skewed towards increased sugar and lipid breakdown minimizes its impact, culminating in overall reduced adult size. Thus, our findings underscore the significant contribution of immune cell metabolic adaptations to adult growth homeostasis, shedding light on the plasticity of this intricate interplay.

The debilitating effects of high sugar on childhood growth trajectory is well established. However, the underlying reasons for how high sugar impairs growth is limited to increased peripheral insulin resistance and resistance to growth hormone signaling. Given animal growth is achieved by complex communication between multiple organs, little is known about how dietary sugar overload impacts the underpinnings of this crosstalk. In the context of immune changes seen in this study, immune-lipid homeostasis has as an adaptive influence on growth where increasing *de novo* lipogenesis can shift the systemic imbalance to drive growth recovery on HSD. In the context of sugar, the sufficiency of immunelactate production in driving growth retardation is significant. The influence of lactacidemia on intrauterine growth retardation (Marconi et al., 1990) concurs with this result and posits immune cell glycolytic shift as a prominent cause for the growth retardation. Thus, we hypothesize that the excessive sugar breakdown with intermediates like lactate shifts the systemic homeostasis on HSD to growth impairment, but when diverted into lipids allows growth on HSD and limits the negative context imposed by sugar metabolism. The induction of *de novo* lipogenesis while allowing an alternate route to metabolize sugars, most likely also restricts pyruvate availability for LDH function, and is perhaps how gain of *ACC* enables growth recovery. The insufficient and inconsistent impact of blocking pyruvate entry into TCA (with loss of PDH) implies the role of TCA in facilitation but not driver of the same. An alternate route that opposes growth more affirmatively is predictably via LDH that raises lactate that then leads to growth impairment on HSD.

The extent of lipogenic induction is limited in immune cells and may be because these cells unlike the fat body are never designed for storage functions. Lipid breakdown is therefore facilitated by bmm and most likely adds to raise the pool of free fatty acids (FFA) and negatively influences growth. FFA and their link with the development of inflammation and insulin insensitivity in peripheral tissues is well established (Johnson and Olefsky, 2013). High levels of circulating FFA and their uptake by non-adipose organs that cannot store fatty acids or their derivatives develop lipotoxicity leading to systemic insulin resistance (Postic and Girard, 2008; Unger, 2003). Thus, it is possible that lactate together with the elevated levels of FFA released from immune cells facilitates growth retardation through invoking insulin resistance. It is also possible that FFA further fuels immune cell TCA activity and adds to growth retardation (Fig. 6g).

In our previous work, we described the role of macrophage internal state as a relevant regulator of animal growth. In this study we extend these observations and show the specifics of internal state changes that are linked to systemic growth homeostasis. Moreover, we found signals that allow immune cells to translate these internal state changes to impact systemic growth. Proand anti-growth factors emanating from immune cells can be perceived by tissues and impinge on a multi-organ cross-talk to define the extent of growth adaption possible. The identification of secreted entities like Hh, that negatively influences growth, while Upd2, Wnt pathway that positively influence growth, illustrate this possibility.

### Macrophages, an important lipogenic organ to ameliorate the effects of dietary sugar induced stress

The protective role of lipogenesis in *Drosophila* and mammals is clearly shown. Lipogenic shift in adipose tissues to store TAGs, protects the animal against sugar induced toxicity (Musselman et al., 2013). Studies in mice have shown that TAG storage protects the animal from developing insulin resistance (Greenberg et al., 2011). Immune cells on the other hand are not conventional storage organs, however, we find that they too respond to HSD exposure in similar ways like the fat body demonstrating a clear lipogenic shift. A transcriptional induction of lipogenic genes with up regulation of TAG synthesis enzymes, *de novo* lipogenesis and alongside requirement for lipid scavenging receptors like Crq, with increasing intracellular levels of TAG, reveal immune cells functioning in the capacity of a key lipogenic organ. The concerted effort of an immune lipogenic state against sugar induced stress, proposes an intriguing role for this circulating organ in enabling processes to accommodate sugar overload. The growth promoting function of this shift further highlights the systemic stress relieving potential enabled by the lipogenic shift in this tissue. Contrary to the known inflammatory consequences of increasing lipid accumulation in immune cells and a defining hallmark of an inflammatory response (Zhang et al., 2022), high sugar induced lipogenic state in immune cells in this study correlates with dampened immunity as evident from the transcriptional down regulation of most immune related pathways.

Specifically, the sugar dependent metabolic reprogramming to drive *de novo* lipogenesis allows commitment of carbon to *de novo* fatty acid synthesis (Parks et al., 2008) and protects the animal against dietary sugar induced stress (Musselman et al., 2013). *de novo* lipogenesis in immune cells also appears to function in a similar capacity. However, the limited expression of *ACC* keeps *de novo* lipogenesis under control. While limited storage of these cell may underlie this restraint, the growth recovery seen with over-expression of *ACC* allude to possible repression on ACC. Given the critical role for ACC at the nexus of metabolism, lipid synthesis and impact on macrophage inflammatory response (Yeudall et al., 2022), it is unclear as to why the immune cells given the overall benefits, would limit their *ACC* expression and consequently their *de novo* lipogenic potential. The possible dysregulation by Hh may underlie this limitation. Hh signaling controls cAMP levels in *Drosophila* larval blood cells (Mondal et al., 2011) and the inhibitory effects of increased cAMP on *de novo* lipogenesis (Batchuluun et al., 2022; von Loeffelholz et al., 2021) aligns well with this possibility but remains to be tested. Alternatively, a limitation in CoA in high sugar conditions (Musselman et al., 2013) may restrict *de novo* lipogenesis or because lipid synthesis via *de novo* lipogenesis is a metabolically costly process compared to the energetically efficient process of direct lipid incorporation into storage forms (Solinas et al., 2015). Therefore, it is very likely that with constant feeding on HSD across the developmental stages, the immune cells resort to a cost-effective process of lipid uptake to maintain a limited lipogenic state.

What is striking is the identification of transcriptional regulators like oxysterol proteins in the screen and their role in promoting growth which remain to be explored in detail. Oxysterols mediate transcriptional regulation that drives direct lipogenesis by promoting lipogenic factors like SREBP and FASN (Horton et al., 2002). Their identification in the screen further strengthens the fact that immune cells are invoked to initiate a lipogenic program. The overall metabolic reshaping exhibited by immune cells to function as a lipogenic organ to confer the tolerance against dietary stress is very striking. Given the specific up regulation of lipogenesis is seen in the face of caloric excess, this is more likely to be a compensatory mechanism to a pathophysiological state, than of a physiological regulation. The data opens a new paradigm to look at these cells in the face of dietary stresses functioning much like fat body or adipose tissues and operating beyond their role in defensive functions.

### Immune cell metabolic heterogeneity: A model to explain the dichotomy of metabolic states seen in HSD

To our knowledge, this is the first study where an unbiased genetic screen and a systematic approach has been conducted to investigate immune interface of animal growth regulation. This extensive analysis has alluded to the existence of unexpectedly contrasting immune metabolic states with opposing consequences on growth phenotype. The dichotomy of immune cell states within the Hml^+^ population is evident across independent and unbiased transcriptomics, genetics and metabolomic approaches.

Macrophage polarization from M1 to M2 state is a well-established concept in mammals (Chen et al, 2023) although not discerned in *Drosophila*, but our data allows us to speculate on a model of immune metabolic heterogeneity that defines immune outcomes. We hypothesize that while all cells may be undergoing metabolic shifts, subpopulations existing in lipogenic or glycolytic state can be proposed and it is a balance in these subtypes that defines the immune outcomes. It is entirely possible that our genetic manipulations introduce bias in immune cells to one type over the other. Recent findings from single cell analysis have alluded to heterogenous population of immune cells in the larvae where lipogenic sub-populations have been identified (Cho et al., 2020; Girard et al., 2021). Moreover, transcriptomics of embryonic versus larval immune cells have revealed distinct metabolic cell states, where embryonic macrophages are lipogenic and glycolytic, while larval immune cells are oxidative, lipolytic and extensively phagocytic (Cattenoz et al., 2020). High sugar induced metabolic reprogramming captures snippets of both larval and embryonic states and therefore the state of heterogeneity could also be an outcome of macrophages from distinct ontogeny contributing to growth. While these are speculations, a balance of proand anti-growth immune states in organismal growth homeostasis is evident from our analysis. Currently limited by bulk transcriptomics studies, single cell analysis of immune cells on HSD will be needed to address the observed immune dichotomy in greater detail.

### Immune cell state at the interface of coordinating systemic organismal growth homeostasis

The indication of macrophages as nutrient sensors (Martínez-Micaelo et al., 2016; Newsholme, 2021) which assimilate environmental information and relay cues at an organismal level is an emerging concept. Evidences supporting immune cells in this capacity have recently been published with respect to governing growth and developmental timing in homeostasis (Juarez-Carreño and Geissmann, 2023; Sriskanthadevan-Pirahas et al., 2023) and in infection (Krejčová et al., 2019). Our findings from the past have elaborated on similar lines (P et al., 2020). They have alluded to systemic influence on insulin signaling and fat body metabolic state and inflammation. Moreover, we have seen that growth recovery on HSD by activating immune-Pvr pathway is independent of insulin activity implying additional players involved in this axis (P et al., 2020). This is where signaling molecules like Adenosine (Bajgar et al., 2015; Bajgar and Dolezal, 2018), Impl2 (Honegger et al., 2008; Krejčová et al., 2023; Okamoto et al, 2013), Upd3 (Romão et al., 2021; Shin et al., 2020; Woodcock et al., 2015), Dilp8 (Colombani et al., 2012; Garelli et al., 2015; Sanchez et al., 2019; Vallejo et al., 2015), Hh (Rodenfels et al., 2014; Vervoort, 2000) and Pvfs (Ghosh et al., 2020; Juarez-Carreño and Geissmann, 2023; Cox et al, 2021) become relevant. Either by modifying insulin resistance or defining developmental timing, they orchestrate metabolic and growth equilibrium. With metabolites like lactate, acetyl CoA and candidate cytokines like Upd2, Wg, Hh and peptide hormones (CCHamide-2, sNPF) identified in the immune cells with specific effect on growth, the ability of these cells to engage in a multiorgan cross-talk beyond insulin signaling can be readily envisaged.

Finally, the concept of altering body size, particularly stunted growth is also considered as an adaptation to adverse environmental conditions rather than a pathological consequence (Bogin et al., 2007). This finds support from studies across humans and other mammals in conditions of heat stress (Elayadeth-Meethal et al., 2018), high altitude (Baye and Hirvonen, 2020; Grant et al., 2022; Pawson and Clegg, 1997) and nutrient availability. In this study, while we focused our discussions on growth recovery in dietary stress, given this is burgeoning problem linked to high sugar diet and childhood growth abnormalities, if any specific contribution these changes have on animal adaptation with respect to survival, fecundity etc., remains to be investigated. Although preliminary, we do observe a benefit on survival on HSD with heightened TCA or glycolytic shift in immune cells. Very strikingly, immune specific downregulation of alpha-ketoglutarate, a key TCA enzyme had heightened larval lethality on HSD (data not shown). It is therefore entirely possible that the limited lipogenic shift, and the stunted growth driven by these pathways is merely an adaptation mechanism to facilitate development in the face of dietary sugar induced stress.

For an animal to grow, it must sense nutrients and then systemically relay information to facilitate nutrient allocation and coordinate metabolism to achieve growth in a wellcoordinated manner. In conditions of nutrient stress, defects in nutrient sensing and signaling is a hallmark feature that is linked to growth abnormalities. Given the physiological complexity of higher model systems, it becomes extremely challenging to address the underlying biological processes and complex crosstalk involved in such responses. The findings in this study provide a fundamental insight into an unexplored and overlooked area of immune metabolic reprogramming and its larger consequences on growth physiology. The small animal sizes observed in macrophage depleted mice and their sensitivity to dietary stresses (Hua et al., 2018; Weisberg et al., 2003) confirms with the larger role played by immune cells in growth control, across systems. Moreover, the connection between immunity and growth (van der Most et al., 2011) alludes to unexplored immune metabolic changes that accompany in these cells in infected conditions (Krejčová et al., 2023) in immune-growth cross-talk. The findings in this study highlights the impact of immune-derived metabolites and the specifics of modulating immune internal metabolic state on decisions defining growth physiology. The cross-talk promoted by immune cells emerges much more than just a consequence of altered immune function and proposes their deterministic role in defining animal growth potential. Further research in this area promises novel insights into growth regulation that currently remain unexplored and beyond our understanding.

## Materials and methods

### Drosophila genetics

Flies were raised on standard cornmeal medium (5% sucrose) at 25°C. For high sugar diet, the sugar content was increased five-fold to 25% sucrose. The RNA*i* lines were obtained either from Bloomington *Drosophila* Stock Center (BDSC, Bloomington, IN) or Vienna *Drosophila* Research Centre (VDRC). The *Gal4* line used was *Hml^Δ^>UAS-GFP* (Sinenko and Mathey-Prevot., 2004) and *w^1118^* flies were used as controls. All genetic crosses were set up at 25°C and then transferred to 29°C where they were grown until analysis either as larvae or as adults. See Table S6 for a complete list of genes and their BDSC or VDRC stock numbers.

### High sugar diet exposure

We utilised two different dietary regimes of high sugar diet (HSD). For the short-term 4hr.HSD regime, *Hml*^△^*>GFP/w^1118^/*RNA*i* feeding third instar larvae (72hr. AEL) reared on regular food (RF, containing 5% sucrose) were transferred to HSD (containing 25% sucrose) where they were allowed to feed for a brief period of four hours only. For Ct.HSD regime, *Hml*^△^*>GFP/w^1118^/*RNA*i* embryos were collected on RF and transferred to HSD (containing 25% sucrose). The larvae were reared at 29°C until feeding 3^rd^ instar stage following which they were processed for experiments related to immune cells and until eclosion for experiments related to adult body size.

### Immune cell counts

For quantification of sessile and circulating immune cells, protocol described by (Petraki et al., 2015) was used to isolate the two immune cell populations. Briefly, three feeding third instar larvae were allowed to bleed for a few seconds in PBS following an incision at both the larval posterior and anterior ends. After the release of the circulating immune cells, the same larvae were transferred to another well and sessile immune cells attached to the larval cuticle released by a process of scrapping and/or jabbing. For quantifying total immune cells, there was no separation of circulating and sessile immune cells. Images were acquired with five fields per sample at 20X magnification. For cell counting, particle analyser in ImageJ was used with size range of 2-infinity. For cell clusters typically counted as one by the software, the number of cells in those clusters were estimated by manual counting. The counting was done for DAPI-positive (representing total blood cells), Hml-positive (Hml^+^), and Hml-negative (Hml^-^) cells. Counting assays were performed in at least two wells per experiment and independently repeated at least three times. The cell numbers obtained were quantified per larva and represented as number of immune cells per mm^2^.

### Immunohistochemistry and staining

For all other experiments except the cell count assay, total immune cells comprising of circulating and sessile pool were analysed. Immune cells were bled and allowed to settle for 30 minutes. Cells were then fixed with 4% formaldehyde in PBS for 10 minutes and incubated with primary antibodies overnight at 4 degrees. Primary antibodies used were rabbit-αACC (1:1000, Jacques Montagne, I2BC, France), mouse-αNICD (1 :50, DSHB, C17.9C6), rabbit-αCi (1 :100, DSHB, 2A1), rabbit-αCrq (1:100, The Scripps Research Institute, La Jolla, USA. The secondary antibody Alexa Fluor 546 (Invitrogen) was used at 1:500. Nuclei were visualized using DAPI (Sigma). Samples were mounted with Vectashield (Vector Laboratories).

For Nile Red staining, formaldehyde fixed cells were incubated in 1:1000 solution of 0.02% Nile Red (Sigma-Cat. No. N3013) for 20 mins, washed and mounted similarly. Images were acquired on Olympus FV3000 confocal microscope with a step size of 1µm at 40X or 60X magnification.

For phalloidin staining, cells were first permeabilized with 0.1% Triton X-100 in PBS (PBST) for 5 mins and then incubated for 2 hours with Atto 565 Phalloidin (Sigma-Aldrich # 94072) diluted 1:100 in PBS. Phalloidin staining was used to assess cell morphology and filipodia length and number. Specifically, for measuring filipodia length, it was done as described in Hao et al, 2018. Briefly, the line tool on Image J was used to draw a line over a filopodium from its tip to cell body with extensions > 0.5 μm being classified as filopodia.

ROS stainings were done as described in (Owusu-Ansah et al., 2008). Larval immune cells were stained with 1:1000 DHE (Dihydroethidium) (Invitrogen, Molecular Probes, D11347) dissolved in 1×PBS for 15 min in the dark. Immune cells were washed in 1×PBS twice and fixed with 4% formaldehyde for 5 mins at room temperature in the dark. After this, 1X PBS wash was given to the immune cells and this step was repeated twice and then Vectashield (Vector Laboratories) was added. The immune cells were imaged immediately.

### Immune cell phagocytosis assay

Immune cells bled in PBS were treated with 0.1µm latex beads (ThermoFischer Scientific #F8801) for 15 minutes and washed three times with PBS to remove the excess free beads. Cells were then fixed with 4% formaldehyde in PBS for 10 minutes, washed with PBS and mounted in Vectashield with DAPI (Vector Laboratories) for imaging. For measuring phagocytic capacity, phagocytic index was measured as number of engulfed latex beads per immune cell (Hao et al, 2018).

### Image analysis and quantification of expression intensities

ImageJ software was used for analysis. For all images, across all experiments, with stainings on circulating immune cells, the quantification of the expression pattern or intensities was done in the following manner. At least two wells per experiment was analysed . Each well had immune cells obtained from a maximum of five larvae. Five to six images were captured for each well at 60X magnification, and the staining was assessed for 5-6 cells/field. The analyses were carried out for at least 60-70 cells per experiment and this was repeated independently at least three times. The quantifications shown in the graphs represents the average expression from these cells across batches. Images were assembled in Adobe Photoshop 2023.

### Immune cell biochemical assays: Triacylglycerol and glucose measurements

TAG and glucose measurements were done as shown in P et al, 2020. Briefly, immune cells bled from at least fifteen larvae per experiment were collected in PBS followed by centrifugation at 1000rpm to pellet the cells.To the pellet, 0.05% 1XPBST (Tween 20) was added and vortexed intermittently by keeping on ice. Protein levels of immune cells were estimated using BCA protein assay kit (ThermoFischer Scientific #23225). For measuring glucose and TAG levels, immune cell samples were first heat inactivated at 70°C for 10 mins and then subjected to metabolite analysis using GOD-POD kit (Sigma#GAGO20) and Triglyceride assay kit (Sigma#T2449) respectively. Assays were performed on Varioskan LUX Multimode Microplate Reader and metabolite levels in each sample were normalised to total protein levels. At least two-three biological replicates were used and the assays were performed in at least three independent experiments (see Legends for “n”, total number of larvae for the assays).

### RNA isolation, bulk RNA-sequencing and Real-Time PCR

Immune cells from thirty-forty feeding 3rd instar larvae fed on RF, 4hr.HSD and Ct.HSD were collected in PBS on ice and stored at -80°C. Total RNA was extracted using Trizol (Life Technologies) followed by assessment of RNA integrity (>7) and purity using an Agilent-2100 Bioanalyzer. Illumina Hi-seq kits were used to construct sequencing libraries following standard protocol, and 100 bp single end reads were generated at Sequencing facility, NCBS (Bengaluru, India).

For Real-Time PCR, RNA was first converted to first-strand cDNA using the SuperScript II Reverse Transcriptase kit (Invitrogen#18064014) following the manufacturer’s instructions. Real-Time PCR was performed in QuantStudio 5 Real-Time PCR System (Applied Biosystems) using SYBR Green Master Mix (Applied Biosystems#A5741) and gene-specific primers. The primers designed using IDT’s Primer Quest Tool are listed in Table S7. Relative quantification of transcript levels was achieved using the Comparative C_t_ method (deltadelta C_t_) using *Rp49* as endogenous control. At least three biological replicates were used and repeated three times (see Legends for “n”, total number of larvae for qPCR)

### RNA-seq data analysis

Post sequencing, 30 to 40 million single-end reads were obtained. FastQC v0.11.5 was used to perform the initial quality check. Adapters were trimmed from the reads using cutadapt v1.8.3 (-a AGATCGGAAGAGCACACGTCTGAACTCCAGTCA). The trimmed reads were mapped to the *Drosophila* genome (*Drosophila melanogaster*. BDGP6.22) using Hisat2 v2.1.0. Read counting was done using featureCounts v2.0.0. DESeq2 v1.40.1 was used to perform the read count normalization and differential expression analysis (Ge et al, 2018; Kanehisa et al, 2000; Kim et al, 2015; Liao et al, 2014; Love et al, 2014; Andrews, S; 2010; Martin, M, 2011). Genes that showed a fold change of at least 2 (up or down), with an adjusted p-value of less than 0.05, were considered as differentially expressed for further analysis. Gene ontology and KEGG pathway enrichment analysis of the differentially expressed genes was done on the ShinyGO v0.60 webserver. Genes that are associated with each metabolic pathway considered here, were retrieved from the KEGG database (http://www.genome.jp/).

### Adult fly size and wing analysis

The adult fly progeny of the tested crosses viz RNA*i* lines with the immune cell specific driver, *Hml*^△>^*GFP* were collected after the eclosion on high sugar diet and kept on 29℃ for one to two days in normal food vials for acclimatization. Both females and males were kept together. After two days the male and female flies were separated, and flies were grouped for imaging for the body size. Each fly was kept in a lateral position and fly wings were moved backward to expose the body. The length from the anterior end of a head to the posterior end of the abdomen in the flies was measured (Lee et al., 2008, 2004). For wingspan quantification, the right wing of each fly was plucked and mounted on a glass slide, separately for males and females. The distilled water was used to mount the wing on a glass slide for proper orientation. The slides were covered with a coverslip and sealed with nail paint. The images of the wings were captured with the Leica MZ 10 F modular stereo microscope and LASX software. Fiji Image J was used to quantify the wing phenotype for wing area using the polygon section tool in the software. The scale was set by converting pixels to millimeters (mms). The hinge region of the wing was excluded while marking the boundary of the wing. More than 50 animals were used for wing span and adult fly length analysis.

### Metabolite extraction and derivatization

For metabolite extraction, blood cells from five feeding 3^rd^ instar (WI) larvae per replicate were extracted and 200μl of 80% ice-cold Methanol was added. After this,100μl of LC/MS grade water was added and the samples were incubated in ice for 30 min. Then 200μl chloroform was added and samples were vortexed for 30 seconds and centrifuged at 13000 RPM for 10 min. at 4°C. The upper phase was transferred into a fresh tube, dried down in a Vacufuge plus speed-vac at room temperature and derivatized further with OB-HA/EDC for metabolite analysis. The interphase was taken for protein estimation for normalization purpose. Proteins were resuspended in 5% SDS and heated at 37°C for 30 minutes. The protein concentration was determined using the Pierce BCA Protein Assay Kit Assay (ThermoFisher). For steady state analysis, the metabolite levels were normalized by per sample per total protein amount in μg.

For derivatization of metabolites (Tan et al., 2014; Walvekar et al., 2018), the dried samples were dissolved in 50 μl of LC/MS grade water and 50μl of 1M EDC (in Pyridine buffer) was added. Samples were kept on a thermomixer for 10 min. at room temperature and 100μl of 0.5 M OBHA (in Pyridine buffer) was added. The samples were incubated again for 1.5 hours on the thermomixer at 25°C, and metabolites were extracted by adding 300 μl of ethyl acetate and this step was repeated three times. Samples were dried down in a Vacufuge plus speed-vac at room temperature and stored at -80°C until run for LC/MS analysis. A minimum of 4 biological replicates were taken per experiment (see Legends for “n”, total number of larvae for metabolomics).

### 13-C labelling and stable isotope tracer analysis

For isotopomer tracer analysis, five feeding 3^rd^ instar (WI) larvae were washed twice in PBS and immune cells were extracted. Blood cells were incubated in 10mM of U^13^C-Pyruvate in 1X PBS (Cambridge Isotope Laboratories, CLM-2440-0.5) for 30 min. Blood-cells were centrifuged down at 13000 RPM for 10 min. and 200ul of 80% ice-cold methanol was added to each sample and stored at -80°C. Samples were further processed for metabolite extraction as done for steady state analysis.

### Liquid chromatography-mass spectrometry (LC/MS) analysis

The metabolite extract was separated using a Waters XBridge C18 Column (2.1 mm, 100 mm, 3.5 mm) coupled to an Agilent QQQ 6470 system. The autosampler and column oven were held at 4°C and 25°C, respectively. The column was used with buffer A (Water and 0.1% Formic Acid) and buffer B (100% acetonitrile and 0.1% Formic Acid). The chromatographic gradient was run at a flow rate of 0.300 ml/minute as follows: 0 min: gradient 10% B; 0.50 min: gradient 10% B; 8 min: gradient 100% B; 10 min: gradient 10% B; 11 min: gradient at 10% B. and 16 min: gradient held at 10%B. The mass spectrometer was operated in MRM, positive ion mode. Mass spectrometry detection was carried out on a QQQ Agilent 6470 system with ESI source. For metabolite quantification, Peak areas were processed using MassHunter workstation (Agilent). Microsoft Excel 2016 and GraphPad Prism 9 software was used for statistical analysis. Q1/Q3 parameters are Retention time (RT) values given in Table S8.

### Sample size and Statistical analyses

In all experiments, n implies the total number of samples analyzed that were obtained from independent experimental repeats and ‘N’ represents the number of independent experimental repeats, which is shown by dot in the graphs and also mentioned in the figure legends. The sample size across all experiments was determined based on the sensitivity of the assay to detect the corresponding changes. All statistical analyses and quantifications were performed using GraphPad Prism Ten and Microsoft Excel 2016. Unpaired t-test with Welch’s correction, Two-way ANOVA with main effects or with Dunnett’s multiple comparison test is employed wherever applicable to account for the variation between and within the experiments (Goyal et al., 2021; Hadjieconomou et al., 2020). P-value given for each graph in legends.

## Data availability

All raw RNAseq reads associated with the study are available from the NCBI SRA (Accession PRJNA1090274).

## Acknowledgements

We thank Prof. Jacques Montagne for the Acetyl-CoA-Carboxylase (ACC) antibody, Prof. Nathalie Franc for the Croquemort (Crq) antibody, the Bloomington *Drosophila* Stock Center (BDSC) and Vienna Drosophila Resource Center (VDRC) for fly stocks. We acknowledge National Centre for Biological Sciences (NCBS), Centre for Cellular and Molecular Platforms (C-CAMP) for Central Imaging & Flow Cytometry Facility (CIFF) and the fly facility. We thank metabolomics facility at MPF, Bindley Bioscience Center, and Dr. Vikki Weake, Purdue University for help with metabolomic experiments. Owing to space limitations, we apologize to our colleagues whose work is not cited. Cartoons were created with BioRender.com. This study was supported by Department of Science and Technology - CRG (Grant number CRG/2021/002815), USIAS Indo French grant awarded to T.M and A.G and DBT/Wellcome Trust Senior Research Fellowship (Grant number IA/S/22/1/506259) awarded to T.M. This work was also supported by the DBT/Wellcome Trust India Alliance Fellowship [Grant number IA/E/18/1/504327] awarded to A.M. S.N is a Graduate Student at inStem, in the Mukherjee lab, and is supported by Council of Scientific & Industrial Research (CSIR)-Fellowship. M.G. is a Graduate Student at inStem, in the Mukherjee lab, and is supported by DST-INSPIRE and, SERB-OVDF fellowship facilitated the work done at Purdue University.

## Supplementary figures legends

**Figure S1.**
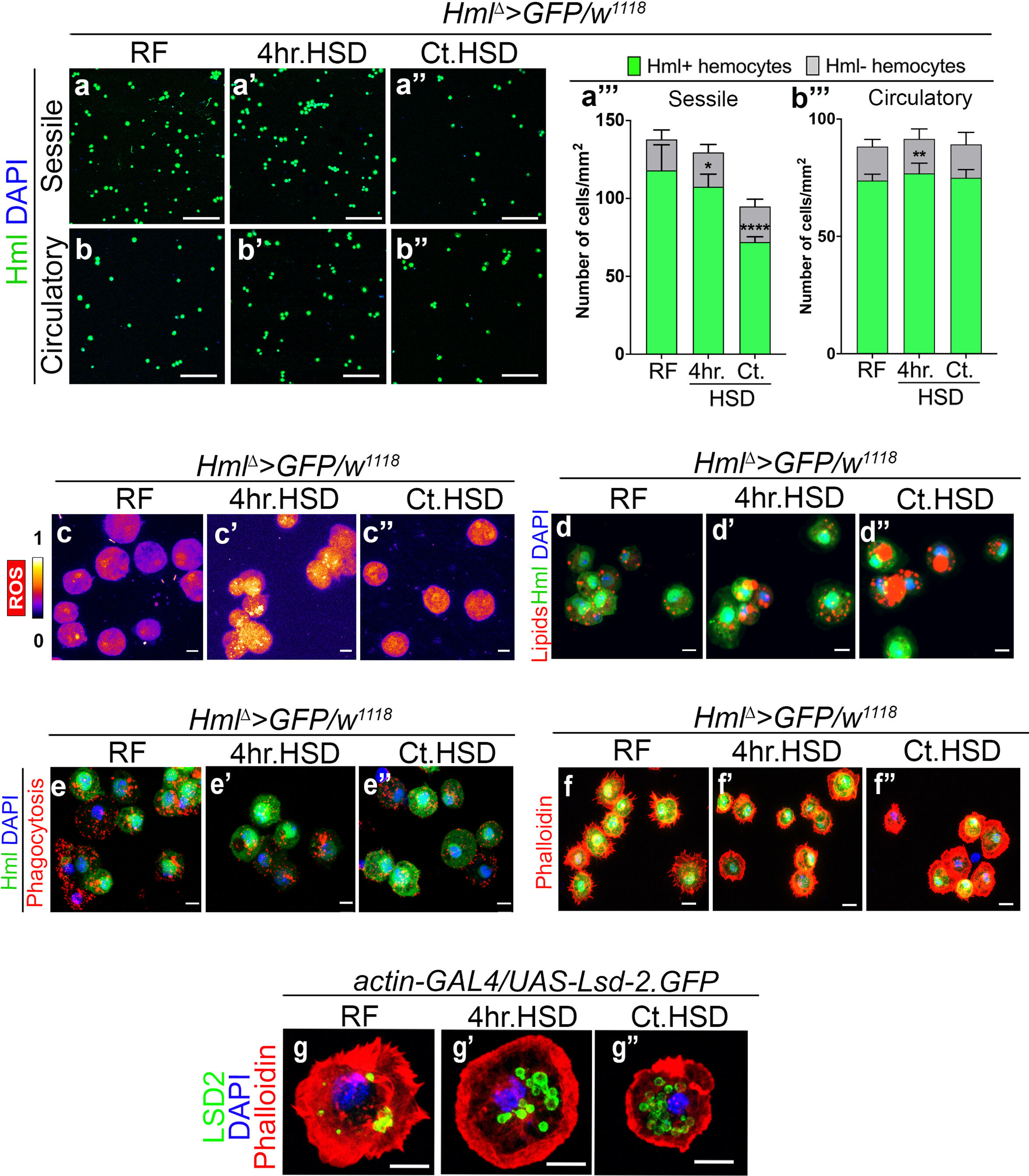
Dietary sugar stress affects larval macrophage physiology. Data information: DNA is stained with DAPI in blue, immune cells are marked in green (*Hml^Δ^>UAS-GFP*). DHE staining to assess ROS, is shown in spectral mode in panels (cc’’). Immune metabolic characterizations using nile red to mark lipids, bead uptake assay to assess phagocytosis, phalloidin to mark actin cytoskeletal changes and *LSD2-GFP* to mark lipid droplets is shown in red in panels (d-g”). Panels (a-b’’), scale bar is 100μm and (c-g’’), scale bar is 5μm. The quantifications in (a’’’) and (b’’’) represent mean with standard deviation. Comparisons for significance are with regular food conditions and asterisks mark statistically significant differences (*p<0.05; **p<0.01; ***p<0.001; ****p<0.0001). The statistical analysis applied is Unpaired t-test with Welch’s correction or Two-way ANOVA with Dunnett’s multiple comparison test wherever applicable. RF, 4hr.HSD and Ct.HSD indicate conditions of larvae fed on regular food (RF), four hours high sugar diet (4hr.HSD) and constitutive high sugar diet (Ct.HSD) respectively. “N” is the total number of experimental repeats and “n” is the total number of larvae analyzed. See methods for further details on larval numbers and sample analysis for each of the experiments. (a-b”’) HSD alters Hml^+^ immune cell numbers. (a-b’’) Representative images of sessile (aa”’) and circulatory immune cells (b-b”’), on RF (a, b), 4hr.HSD (a’, b’) and Ct.HSD (a”, b”). Compared to sessile immune cells in (a) *Hml^Δ^>GFP/w^1118^*(*Control* (RF), (a’) *Hml^Δ^>GFP/w^1118^* (4hr.HSD) did not show any dramatic change in their numbers, but (a’’) *Hml^Δ^>GFP/w^1118^* (Ct.HSD) larvae show significant reduction in Hml^+^ sessile population. See representative quantifications in a”’. Circulating cell numbers in (b’) *Hml^Δ^>GFP/w^1118^* (4hr.HSD) and (b’’) *Hml^Δ^>GFP/w^1118^* (Ct.HSD) showed no striking difference compared to the *Control* (b). See representative quantifications in b”’. (a’’’) Quantification of sessile Hml^+^ immune cell numbers in *Hml^Δ^>GFP/w^1118^* (*Control,* RF, N=3, n=18), *Hml^Δ^>GFP/w^1118^* (4hr.HSD, N=3, n=18, p=0.0107) and *Hml^Δ^>GFP/w^1118^* (Ct.HSD, N=3, n=18, p<0.0001). (b’’’) Quantification of circulatory Hml^+^ immune cell numbers in *Hml^Δ^>GFP/w^1118^* (*Control,* RF, N=3, n=18), *Hml^Δ^>GFP/w^1118^* (4hr.HSD, N=3, n=18, p=0.0072) and *Hml^Δ^>GFP/w^1118^* (Ct.HSD, N=3, n=18, p=0.1996). (c-c’’) Representative images of immune cells to assess ROS levels. (c) ROS level in immune cell of *Hml^Δ^>GFP/w^1118^* (*Control*, RF). (c’) *Hml^Δ^>GFP/w^1118^* (4hr.HSD) and (c’’) *Hml^Δ^>GFP/w^1118^* (Ct.HSD) show increased ROS levels as compared to *Control* (c). (d-d’’) Representative images of immune cells with Nile red staining to assess for lipid droplet accumulation. (d) *Hml^Δ^>GFP/w^1118^*(*Control,* RF). (d’) *Hml^Δ^>GFP/w^1118^* (4hr.HSD) and (d’’) *Hml^Δ^>GFP/w^1118^* (Ct.HSD) show gradual increase in immune cell lipid content compared to *Control* (d). (e-e’’) Representative confocal images of immune cells to assess phagocytosis through bead uptake assay 15 min post exposure. (e) *Hml^Δ^>GFP/w^1118^*(*Control*, RF). (e’) *Hml^Δ^>GFP/w^1118^* (4hr.HSD) and (e’’) *Hml^Δ^>GFP/w^1118^* (Ct.HSD) show reduction in number of internalised beads when compared to *Control* (e). (f-f’’) Representative confocal images of immune cells assessed for cellular morphology, (f) *Hml^Δ^>GFP/w^1118^* (*Control*, RF). (f’) *Hml^Δ^>GFP/w^1118^* (4hr.HSD) and (f’’) *Hml^Δ^>GFP/w^1118^*(Ct.HSD) show reduction both in number as well as in length of filopodia compared to *Control* (f). (g-g’’) Representative images of immune cells assessed for lipid droplets with *UAS-LSD2-GFP* reporter line. (g) *actin-GAL4/UAS-LSD2-GFP* (*Control,* RF). (g’) *actin-GAL4/UAS-LSD2-GFP* (4hr.HSD) and (g’’) *actin-GAL4/UAS-LSD2-GFP* (Ct.HSD) show gradual increase in immune cell lipid droplets (green) compared to *Control* (g).

**Figure S2.**
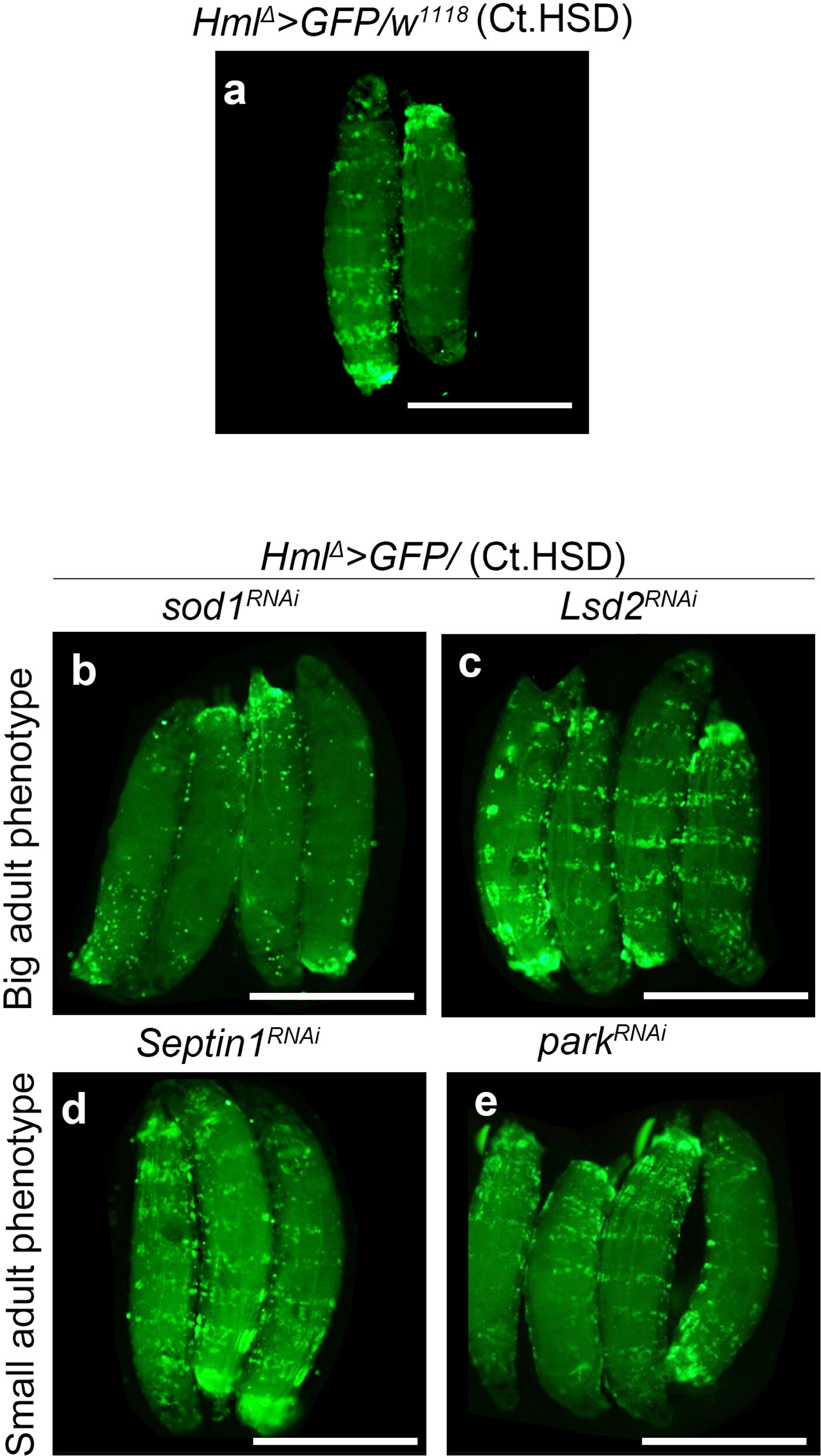
Immune cell state rather than numbers determines growth on HSD. (a-e) Representative third instar larval images of a few candidate genes from the genetic screen. (a) *Hml^Δ^>GFP/w^1118^* (*Control*, Ct.HSD). Big fly size is seen in (b) *Hml^Δ^>GFP/Sod1^RNAi^* and (c) (*Hml^Δ^>GFP/Lsd2^RNAi^).* Small fly size is seen in (d) (*Hml^Δ^>GFP/ Septin1^RNAi^*) and (e) (*Hml^Δ^>GFP/park^RNAi^*). Knock down of the genes in immune cells and the corresponding change on adult fly size allude to immune cell state and its important contribution on growth, as opposed to their numbers.

**Figure S3.**
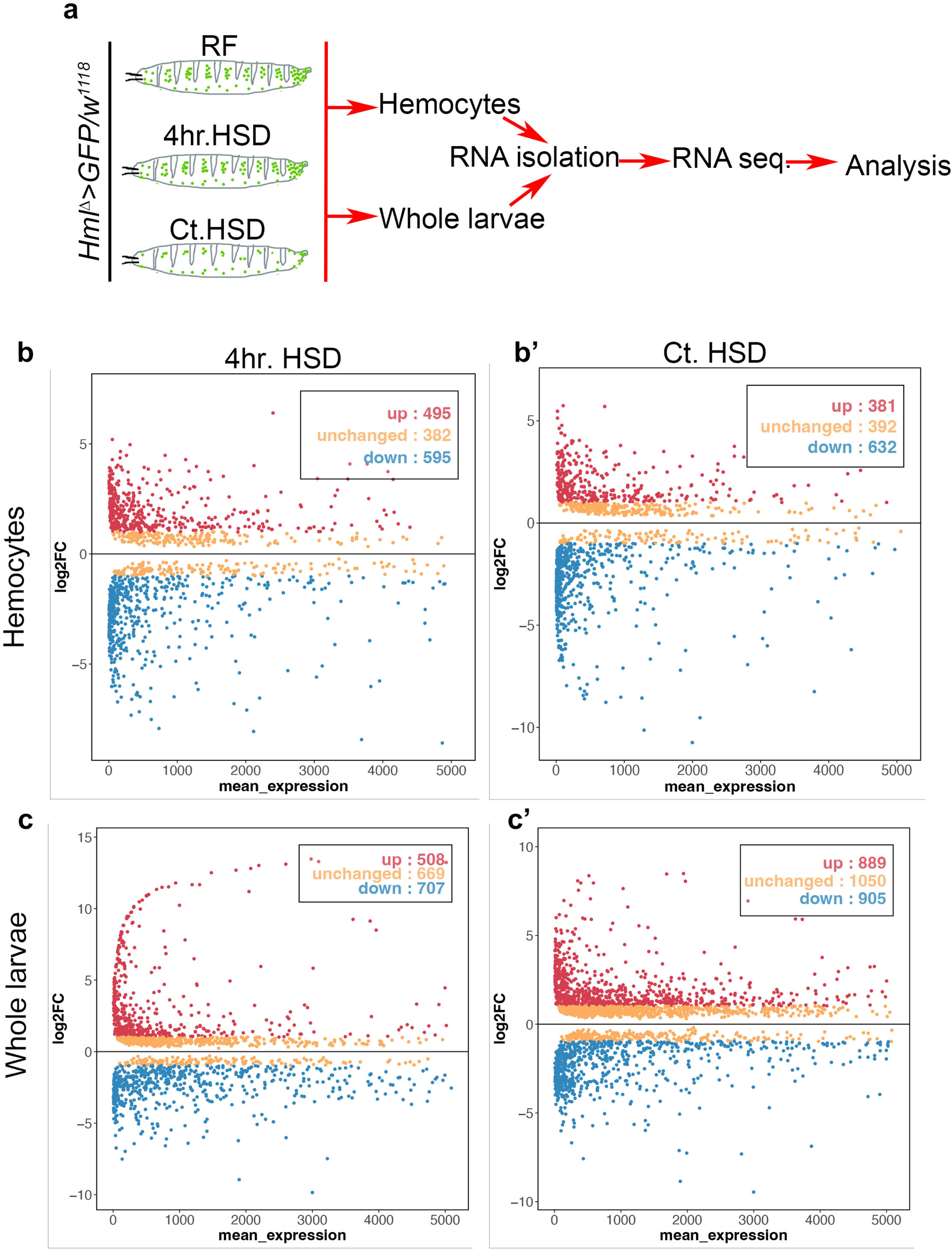
Whole-genome transcriptomics of immune cells and larvae exposed to high sugar diet. (a) Schematic representation of transcriptomics performed on immune cells and whole larvae in *Hml^Δ^GFP>/w^1118^* (Control, RF), *Hml^Δ^GFP>/w^1118^*(4hr.HSD) and *Hml^Δ^GFP>/w^1118^* (Ct.HSD) dietary conditions. See methods for details. (b-b’) Scatter plots depicting the distribution of differentially expressed genes in immune cells and (c-c’) whole larvae fed on 4hr.HSD and Ct.HSD respectively compared to larvae fed on RF.

**Figure S4.**
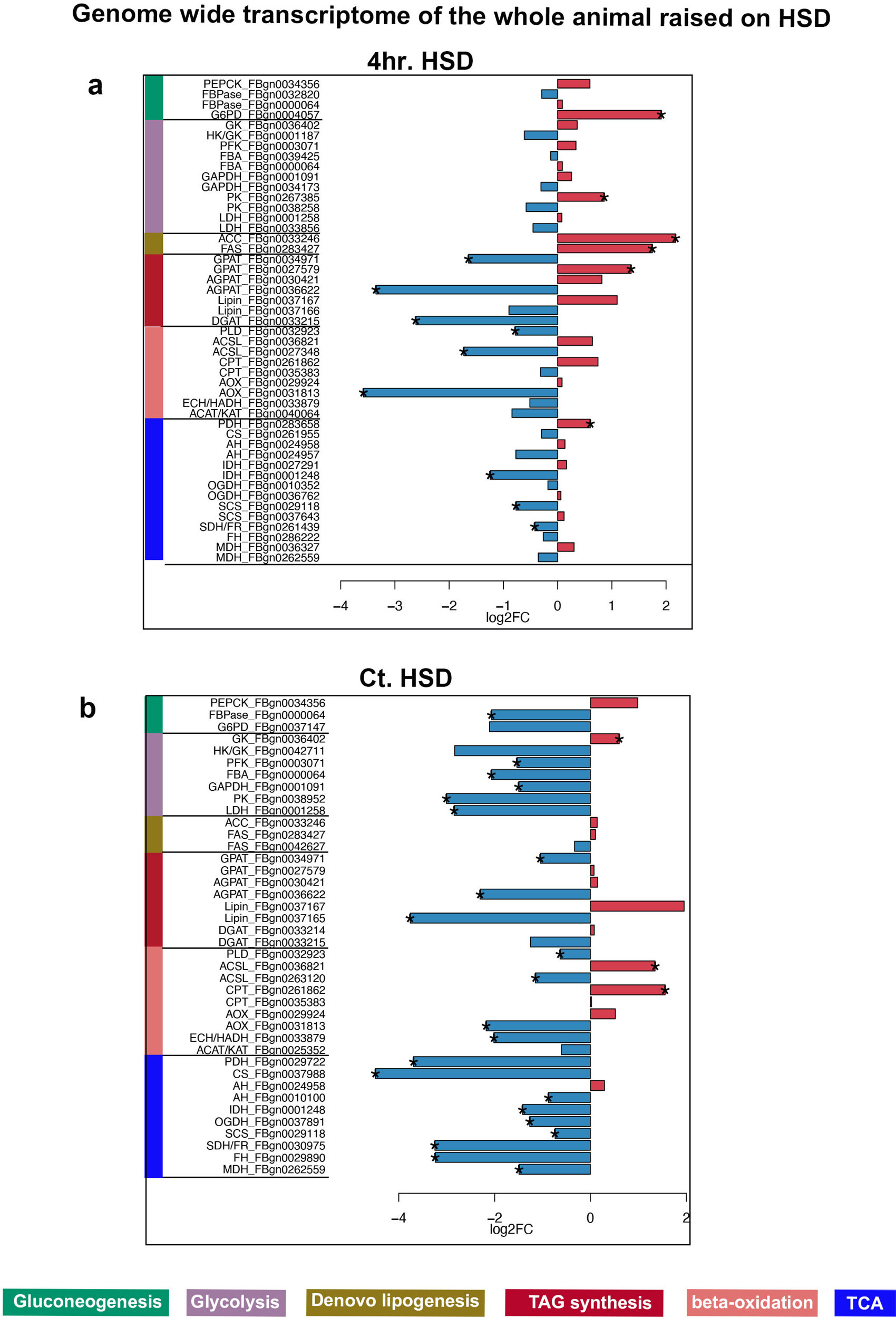
High sugar diet dampens metabolic events in whole larvae. (a) and (b) Bar plots of up regulated (in red) and down regulated genes (in blue) of different metabolic pathways in whole animal (larvae) raised on 4hr.HSD and Ct.HSD respectively. Metabolic genes are down-regulated in 4hr.HSD whole larvae and this is sustained in long term Ct.HSD animals.

**Figure S5.**
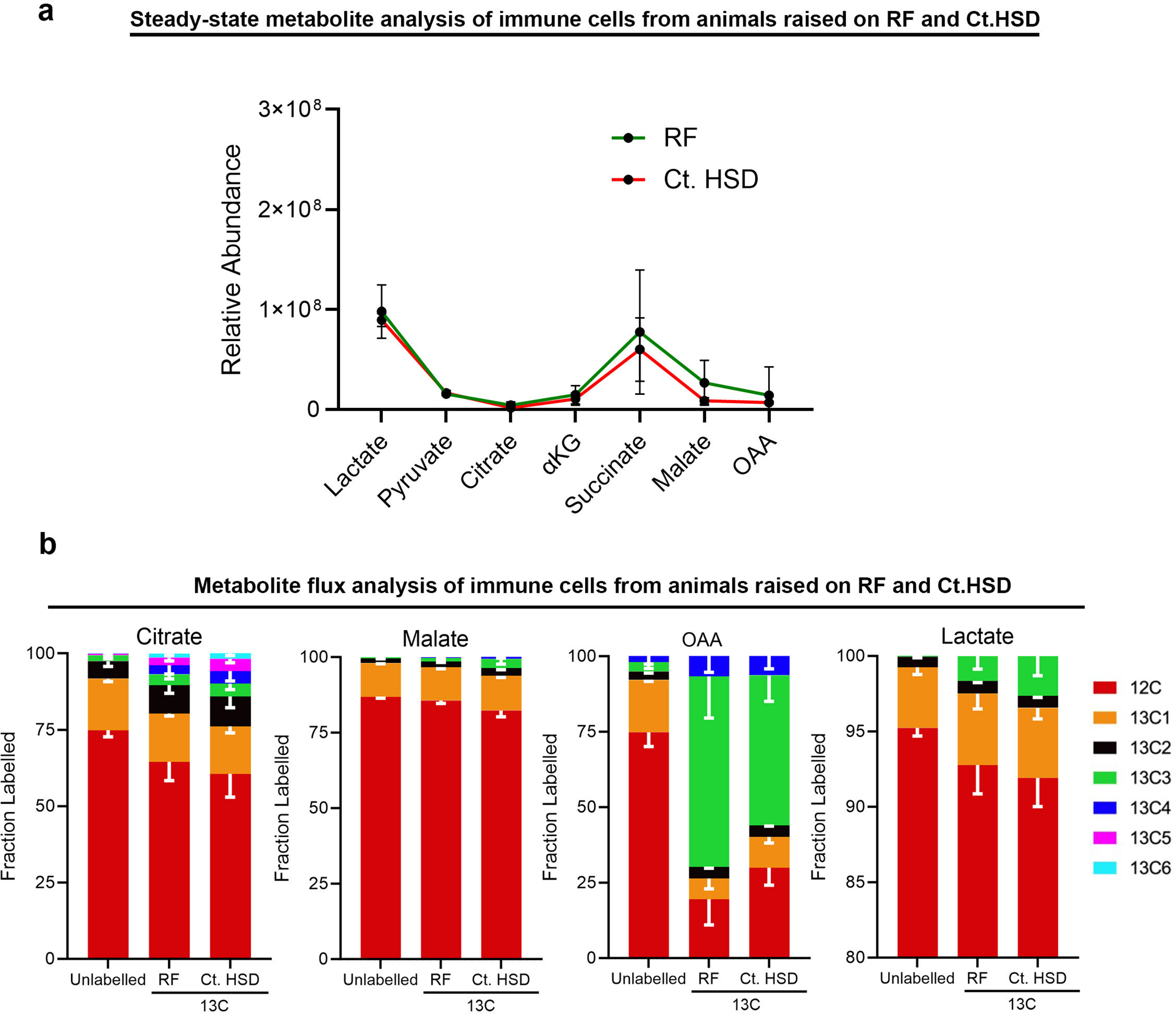
Steady-state metabolite and flux analysis with U^13^C-pyruvate from immune cells on RF and Ct.HSD. (a) Mass spectrometry analysis of steady-state lactate and TCA metabolites in immune cells between *Hml^Δ^GFP>/w^1118^* (Control, RF) and *Hml^Δ^GFP>/w^1118^* (Ct.HSD) do not show any significant change. (b) Distribution of labeled U^13^C pyruvate in TCA metabolites and lactate in *Hml^Δ^GFP>/w^1118^* (Control, RF) and *Hml^Δ^GFP>/w^1118^* (Ct.HSD) conditions showing the respective fraction label incorporation from 13C. 13C label incorporation in unlabelled condition is shown to indicate the natural isotopic abundance. Ct.HSD led to an increase in M+5 label incorporation in citrate, label incorporation in malate M+2 and a decrease in M+3 label incorporation in OAA. M+3 label incorporation in lactate increases in Ct.HSD condition.

**Figure S6.**
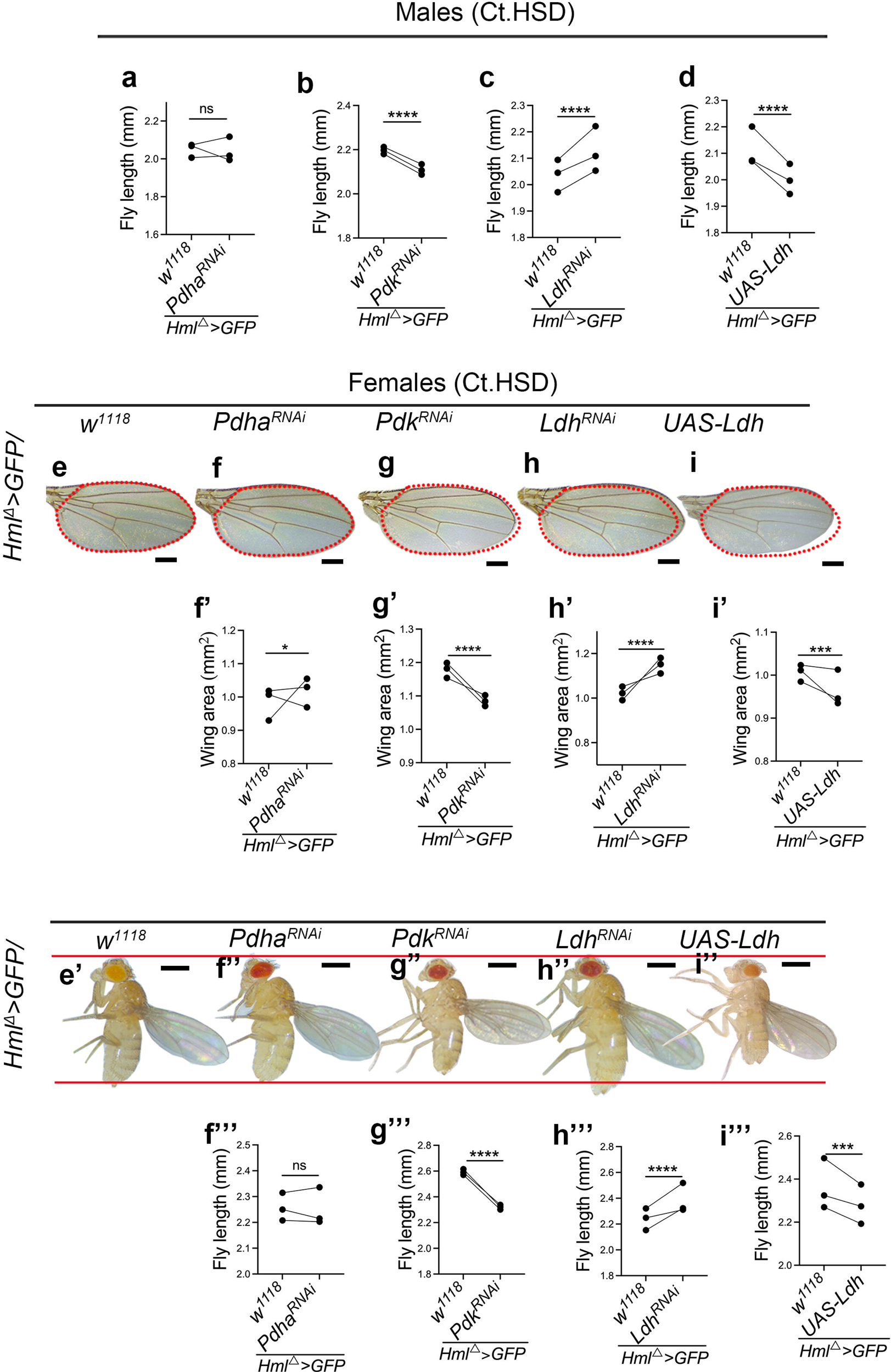
An oxidative and aerobic glycolytic state in immune cells represses growth on HSD. Data information: Scale bar: 0.5mm for flies and 0.25mm for wings. In quantification graphs, shown in panel (a-d), (f’-i’), (f’’’-i’’’), each dot represents an experimental repeat. Comparison for significance is with respect to Control on HSD. Asterisks mark statistically significant differences (*p<0.05; **p<0.01; ***p<0.001; ****p<0.0001). The statistical analysis applied is Two-way ANOVA, main effects test. “N” is the total number of repeats. “n” is the total number of animals analyzed. Only right wing from each adult fly was selected for quantification. The red dotted line in the panels marks the wing span area and is used as a reference to showcase any change in wing span area across genotypes. The two horizontal red lines in the panels is used as a reference to showcase any change in fly lengths across genotypes. RF and Ct.HSD correspond to regular food and constitutive high sugar diet respectively. (a-i’’’) Modulating larval immune cell TCA and glycolytic activity affects adult growth. (a-d) Fly length quantifications in males. (a) *Hml^Δ^>GFP/Pdha^RNAi^*(N=3, n=65, p=0.8628, in comparison to corresponding Ct.HSD *Control*, *Hml^Δ^>GFP/w^1118,^* N=3, n=42), (b) *Hml^Δ^>GFP/Pdk^RNAi^*(N=3, n=77, p<0.0001 in comparison to corresponding Ct. HSD *Control*, *Hml^Δ^>GFP/w^1118,^* N=3, n=62), (c) *Hml^Δ^>GFP/Ldh^RNAi^*(N=3, n=87, p<0.0001 in comparison to Ct.HSD *Control*, *Hml^Δ^>GFP/w^1118,^* N=3, n=65) and (d) *Hml^Δ^>GFP/UAS-Ldh* (N=3, n=83, p<0.0001 in comparison to Ct.HSD *Control*, *Hml^Δ^>GFP/w^1118,^* N=3, n=71). Representative images of wings of adult females (e-i) showing size phenotype on Ct.HSD from respective genetic backgrounds. Compared to (e) Ct.HSD *Control* (*Hml^Δ^>GFP/w^1118^*), (f) expressing *Pdha^RNAi^* (*Hml^Δ^>GFP/Pdha^RNAi^*) in immune cells to reduce TCA activity did not show any striking change in wing span. Expressing (g) *Pdk^RNAi^* (*Hml^Δ^>GFP/Pdk^RNAi^*) to increase TCA activity, contrarily decreases size. (h) Down regulating immune cell glycolytic activity by expressing *Ldh^RNAi^* (*Hml^Δ^>GFP/Ldh^RNAi^*) causes increase in size and (i) further increasing *Ldh* expression (*Hml^Δ^>GFP/UAS-Ldh*) led to decrease in size. (f’-i’) Wing span quantifications in females. (f’) *Hml^Δ^>GFP/Pdha^RNAi^*(N=3, n=57, p=0.0481 in comparison to corresponding Ct.HSD *Control*, *Hml^Δ^>GFP/w^1118,^*N=3, n=67), (g’) *Hml^Δ^>GFP/Pdk^RNAi^* (N=3, n=79, p<0.0001 in comparison to corresponding Ct.HSD *Control*, *Hml^Δ^>GFP/w^1118,^*N=3, n=61), (h’) *Hml^Δ^>GFP/Ldh^RNAi^* (N=3, n=106, p<0.0001 in comparison to corresponding Ct.HSD *Control*, *Hml^Δ^>GFP/w^1118,^*N=3, n=90) and (i’) *Hml^Δ^>GFP/UAS-Ldh* (N=3, n=104, p=0.0005 in comparison to corresponding Ct.HSD *Control*, *Hml^Δ^>GFP/w^1118,^* N=3, n=81). (f’’-i’’) Representative images of adult females on Ct.HSD from respective genetic backgrounds compared to control (e’). (f’’’-i’’’) Fly length quantifications in females. (f’’’) *Hml^Δ^>GFP/Pdha^RNAi^*(N=3, n=60, p=0.7817 in comparison to corresponding Ct.HSD *Control*, *Hml^Δ^>GFP/w^1118,^* N=3, n=73), (g’’’) *Hml^Δ^>GFP/Pdk^RNAi^* (N=3, n=81, p<0.0001 in comparison to corresponding Ct.HSD *Control*, *Hml^Δ^>GFP/w^1118,^*N=3, n=62), (h’’’) *Hml^Δ^>GFP/Ldh^RNAi^* (N=3, n=70, p<0.0001 in comparison to Ct.HSD *Control*, *Hml^Δ^>GFP/w^1118,^* N=3, n=58) and (i’’’) *Hml^Δ^>GFP/UAS-Ldh* (N=3, n=84, p=0.0007 in comparison to Ct.HSD *Control*, *Hml^Δ^>GFP/w^1118,^* N=3, n=71).

**Figure S7.**
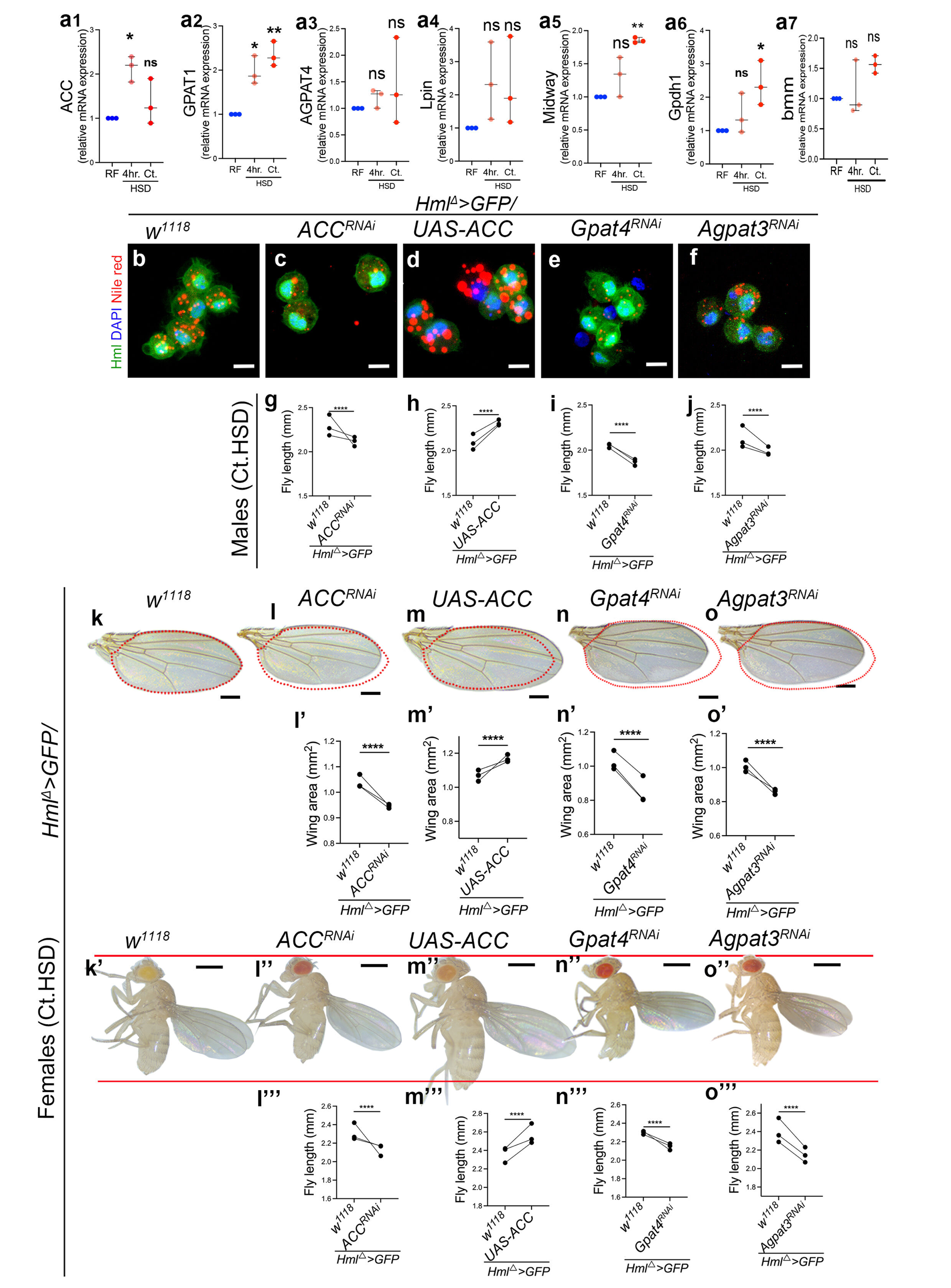
Immune cell lipid homeostasis and systemic growth regulation on HSD. Data information: DNA is stained with DAPI (blue), immune cells are shown in green (*Hml^Δ^>UAS-GFP*). Nile red staining to mark lipids is shown in red in panels (b-f). Scale bar: 5μm for immune cells, 0.5mm for flies and 0.25mm for wings. In quantification graphs (a1-a7), (g-j), (l’-o’) and (l’’’-o’’’), each dot represents an experimental repeat. Except for panel (a1-a7) and (b-f), where comparisons are with respect to Control on RF, in all other panels comparison for significance is with respect to Control on HSD. Asterisks mark statistically significant differences (*p<0.05; **p<0.01; ***p<0.001; ****p<0.0001). The statistical analysis applied is Two-way ANOVA, main effects or with Dunnett’s multiple comparison test wherever applicable. “N” is the total number of repeats. “n” is the total number of animals (larvae and adult flies). Only right wing from each adult fly was selected for quantification. The red dotted line in the panels marks the wing span area and is used as a reference to showcase any change in wing span area across genotypes. The two horizontal red lines in the panels is used as a reference to showcase any change in fly lengths across genotypes. RF, 4hr.HSD and Ct.HSD correspond to regular food, four hours high sugar diet and constitutive high sugar diet respectively. See methods for further details on larval numbers and sample analysis for each of the experiments. (a1-a7) Relative immune-specific expression of lipid metabolism genes by Real-Time PCR show; (a1) a significant up regulation of *ACC*, at 4hr.HSD (*Hml^Δ^>GFP/w^1118,^* N=3, n=105, p=0.0167) but not with Ct.HSD (*Hml^Δ^>GFP/w^1118,^* N=3, n=120, p=0.3690). (a2) *GPAT1* (N=3, n=120, p=0.0044) was up regulated in Ct.HSD. (a3) *AGPAT4* and (a4) *Lpin* did not show any change. (a5) *Midway* (N=3, n=120, p=0.0052) and (a6) *Gpdh1* (N=3, n=120, p=0.0165) were up regulated in Ct.HSD and (a7) *bmm* did not show any change at 4hr.HSD (N=3, n=105) and Ct.HSD (N=3, n=120). All comparisons are with control *Hml^Δ^GFP>/w^1118^*on RF (N=3, n=105). (b-f) Representative images of immune cells stained to observe lipid droplets (Nile red, red) in (b) Ct.HSD Control (*Hml^Δ^>GFP/w^1118^*) and with immune-specific (c) loss of *ACC* function, (*Hml^Δ^>GFP/ACC^RNAi^*), (d) gain of *ACC* expression (*Hml^Δ^>GFP/UAS-ACC*), (e) loss of *Gpat4* (*Hml^Δ^>GFP/Gpat4^RNAi^*) and (f) loss of *Agpat3* (*Hml^Δ^>GFP/Agpat3^RNAi^)* function. Compared to Ct.HSD Control (b), loss of immune cells lipid synthesis both *de novo* (c) or TAG synthesis (e) and (f) led to reduced lipid droplets in them. Contrarily, gain of *de novo* lipid synthesis (d) shows increased lipid droplets in them. (g-o’’’) Modulating larval immune cell lipid homeostasis affects adult growth. (g-j) Fly length quantifications in males. (g) *Hml^Δ^>GFP/ACC^RNAi^*(N=3, n=44, p<0.0001) compared to corresponding Ct.HSD *Control*, (*Hml^Δ^>GFP/w^1118,^*N=3, n=45), (h) *Hml^Δ^>GFP/UAS-ACC* (N=3, n=43, p<0.0001) compared to corresponding Ct.HSD *Control*, (*Hml^Δ^>GFP/w^1118,^* N=3, n=56), (i) *Hml^Δ^>GFP/Gpat4^RNAi^*(N=3, n=90, p<0.0001) compared to Ct.HSD *Control*, (*Hml^Δ^>GFP/w^1118,^*N=3, n=75) and (j) *Hml^Δ^>GFP/Agpat3^RNAi^* (N=3, n=80, p<0.0001) compared to Ct.HSD *Control*, (*Hml^Δ^>GFP/w^1118,^* N=3, n=78). (k-o) Representative images of wings of adult females showing size phenotype on Ct.HSD from respective genetic backgrounds. Compared to (k) Ct.HSD *Control* (*Hml^Δ^>GFP/w^1118^*), (l) loss of *ACC* function (*Hml^Δ^>GFP/ACC^RNAi^*) leads to growth retardation while (m) gain of immune *ACC* expression (*Hml^Δ^>GFP/UAS-ACC*) shows growth recovery and the flies are much larger than Ct.HSD *Control* adults (k). Similarly, loss of TAG synthesis, by blocking (n) *Gpat4* (*Hml^Δ^>GFP/Gpat4^RNAi^*) or (o) *Agpat3* (*Hml^Δ^>GFP/Agpat3^RNAi^*) shows reduction in animal size. (l’-o’) Wing span quantifications in females. (l’) *Hml^Δ^>GFP/ACC^RNAi^*(N=3, n=84, p<0.0001) compared to Ct.HSD *Control*, (*Hml^Δ^>GFP/w^1118,^*N=3, n=115), (m’) *Hml^Δ^>GFP/UAS-ACC* (N=3, n=83, p<0.0001) compared to Ct.HSD *Control*, (*Hml^Δ^>GFP/w^1118^*, N=3, n=106), (n’) *Hml^Δ^>GFP/Gpat4^RNAi^* (N=3, n=96, p<0.0001) compared to Ct.HSD *Control*, (*Hml^Δ^>GFP/w^1118^*, N=3, n=88) and (o’) *Hml^Δ^>GFP/Agpat3^RNAi^* (N=3, n=42, p<0.0001) compared to Ct.HSD *Control*, (*Hml^Δ^>GFP/w^1118^*, N=3, n=61). (l’’-o’’) Representative images of adult females on Ct.HSD from respective genetic backgrounds compared to control (k’). (l’’’-o’’’) Fly length quantifications in females. (l’’’) *Hml^Δ^>GFP/ACC^RNAi^*(N=3, n=33, p<0.0001) compared to corresponding Ct.HSD *Control*, (*Hml^Δ^>GFP/w^1118,^* N=3, n=40), (m’’’) *Hml^Δ^>GFP/UAS-ACC* (N=3, n=58, p<0.0001) compared to corresponding Ct.HSD *Control*, (*Hml^Δ^>GFP/w^1118,^*N=3, n=45), (n’’’) *Hml^Δ^>GFP/Gpat4^RNAi^* (N=3, n=78, p<0.0001) compared to Ct.HSD *Control*, (*Hml^Δ^>GFP/w^1118,^* N=3, n=78) and (o’’’) *Hml^Δ^>GFP/Agpat3^RNAi^* (N=3, n=56, p<0.0001) compared to Ct.HSD *Control*, (*Hml^Δ^>GFP/w^1118,^* N=3, n=73).

**Figure S8.**
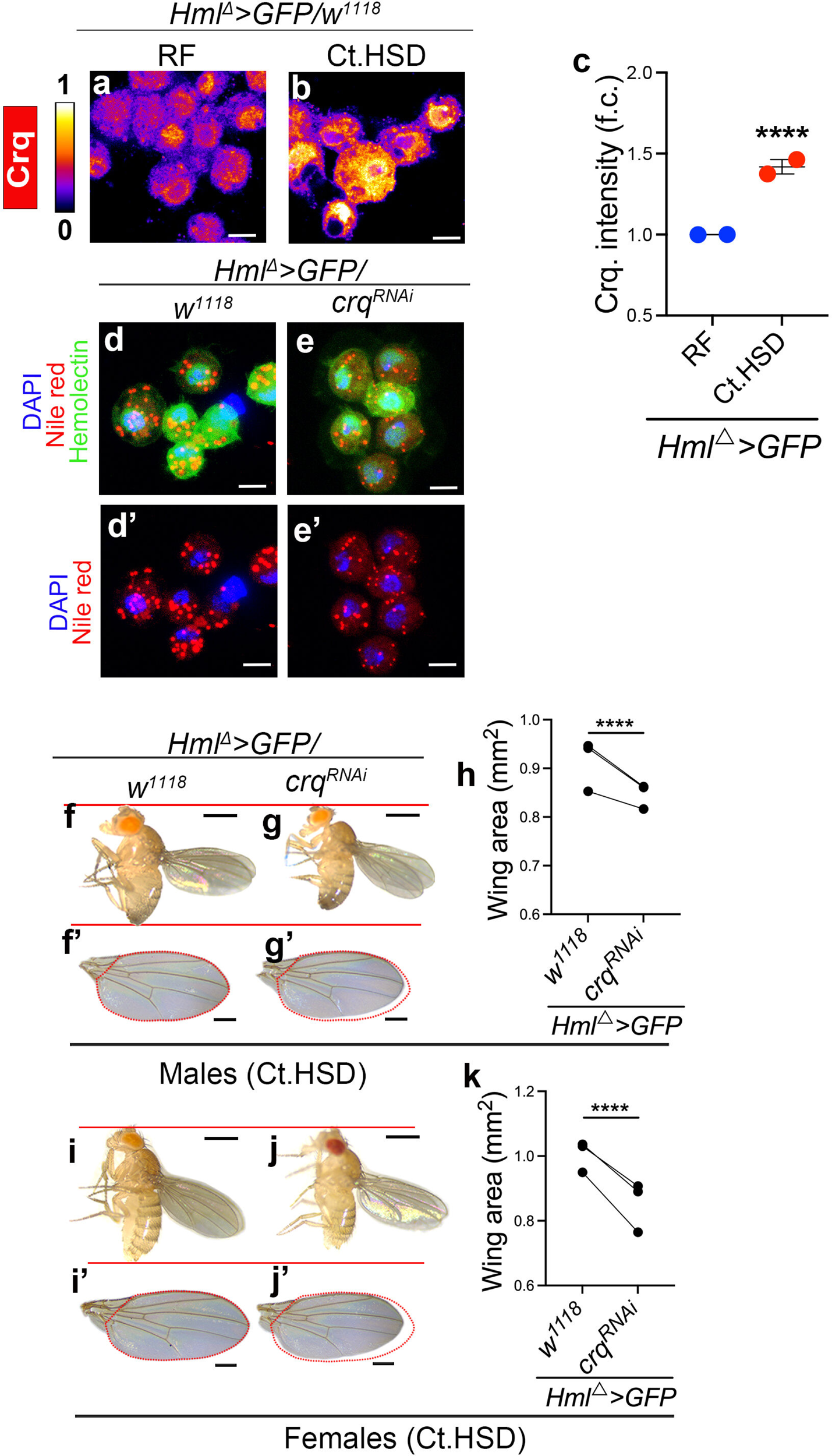
Immune cell lipid uptake regulates growth on high sugar diet. Data information: DNA is stained with DAPI (blue), immune cells are shown in green (*Hml^Δ^>UAS-GFP*). Croquemort (Crq) staining is shown in spectral mode in panels (a-b). Nile red staining to mark lipids is shown in red in panels (d-e’). Scale bar: 5μm for immune cells, 0.5mm for flies and 0.25mm for wings. In quantification graphs (c), (h), and (k), each dot represents an experimental repeat. Except for panel (a-c) where comparisons are with respect to Control on RF, in all other panels comparison for significance is with respect to Control on HSD. Asterisks mark statistically significant differences (*p<0.05; **p<0.01; ***p<0.001; ****p<0.0001). The statistical analysis applied is Two-way ANOVA, main effects. “N” is the total number of repeats. “n” is the total number of animals (larvae and adult flies). Only right wing from each adult fly was selected for quantification. The red dotted line in the panels marks the wing span area and is used as a reference to showcase any change in wing span area across genotypes. The two horizontal red lines in the panels is used as a reference to showcase any change in fly lengths across genotypes. RF and Ct.HSD correspond to regular food and constitutive high sugar diet respectively. See methods for further details on larval numbers and sample analysis for each of the experiments. (a-b) Representative images of immune cells stained to visualize Crq protein expression. Compared to Crq protein levels in (a) RF Control (*Hml^Δ^>GFP/w^1118^*), (b) long-term HSD condition (Ct.HSD) the expression of Crq is increased dramatically. (c) Relative quantification of Crq protein expression, RF (N=3, n=30) and Ct.HSD (N=2, n=30, P<0.0001). (d-e’) Representative images of immune cells on Ct.HSD stained to show lipid droplets (Nile Red, red). Compared to lipid levels seen in (d, d’) HSD *Control* (*Hml^Δ^>GFP/w^1118^*), (e, e’) loss of immune cell *crq* function (*Hml^Δ^>GFP/crq^RNAi^*) leads to decrease in lipid droplets. (f-g’) Representative images of (f-g) adult males and (f’-g’) right-wing to show size phenotype on Ct.HSD upon manipulating lipid uptake in immune cells. Compared to (f-f’) HSD *Control* (*Hml^Δ^>GFP/w^1118^*), (g-g’) loss of *crq* (*Hml^Δ^>GFP/crq^RNAi^*) causes a further reduction in body size. (h) Male wingspan quantification of Ct.HSD control (*Hml^Δ^GFP>/w^1118^*, N=3, n=124), and *Crq^RNAi^* (*Hml^Δ^GFP>/Crq^RNAi^,* N=3, n=51, p<0.0001). (i-j’’) Representative images of (i-j) adult females and (i’-j’) wing to show size phenotype on Ct.HSD upon manipulating lipid uptake in immune cells. Compared to (i-i’) Ct.HSD *Control* (*Hml^Δ^>GFP/w^1118^*), (jj’) loss of *crq* (*Hml^Δ^>GFP/crq^RNAi^*) causes a further reduction in body size. (k) Female wingspan quantification of Ct.HSD control (*Hml^Δ^GFP>/w^1118^*, N=3, n=111), and c*rq^RNAi^* (*Hml^Δ^GFP>/crq^RNAi^,*N=3, n=31, p<0.0001).

**Figure S9.**
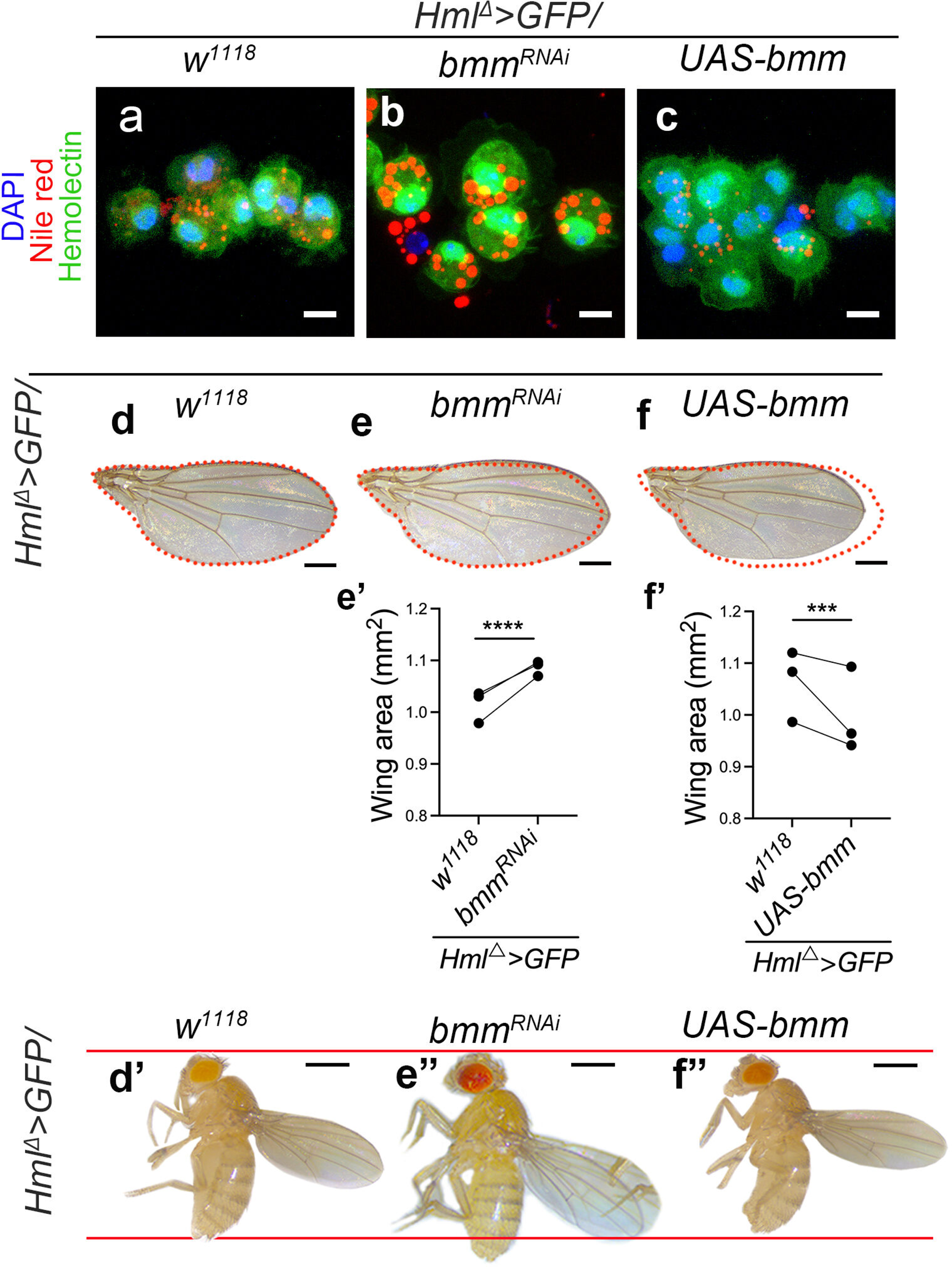
Immune cell lipolytic state as inhibitor of systemic growth on HSD. Data information: DNA is stained with DAPI (blue), immune cells are shown in green (*Hml^Δ^>UAS-GFP*). Nile red staining to mark lipids is shown in red in panels (a-c). Scale bar: 5μm for immune cells, 0.5mm for flies and 0.25mm for wings. In quantification graphs e’-f’), each dot represents an experimental repeat. Comparison for significance is with respect to Control on HSD. Asterisks mark statistically significant differences (*p<0.05; **p<0.01; ***p<0.001; ****p<0.0001). The statistical analysis applied is Two-way ANOVA, main effects. “N” is the total number of repeats. “n” is the total number of animals (larvae and adult flies). Only right wing from each adult fly was selected for quantification. The red dotted line in the panels marks the wing span area and is used as a reference to showcase any change in wing span area across genotypes. The two horizontal red lines in the panels is used as a reference to showcase any change in fly lengths across genotypes. RF and Ct.HSD correspond to regular food and constitutive high sugar diet respectively. (a-c) Representative images of immune cells on Ct.HSD stained to show lipid droplets (Nile Red, red). Compared to lipid levels seen in (b) Ct.HSD *Control* (*Hml^Δ^>GFP/w^1118^*), (c) loss of immune cell *brummer* function (*Hml^Δ^>GFP/bmm^RNAi^*) led to increase in lipid droplets and (d) gain of *bmm* expression (*Hml^Δ^>GFP/UAS-bmm*) led to decrease in lipids respectively. (d-f’’) Modulating larval immune cell lipolysis affects adult growth. Representative images of fly wings of adult females (d-f) showing size phenotype on Ct.HSD from respective genetic backgrounds. Compared to (d) Ct.HSD *Control* (*Hml^Δ^>GFP/w^1118^*), (e) loss of *bmm* (*Hml^Δ^>GFP/bmm^RNAi^*) or (f) increase in its expression (*Hml^Δ^>GFP/UAS-bmm*) in immune cells causes either a recovery in adult fly size or a further reduction in size respectively. (e’-f’) Quantification of female wingspan in (e’) *Hml^Δ^>GFP/bmm^RNAi^*(N=3, n=108, p<0.0001) in comparison to HSD *Control*, (*Hml^Δ^>GFP/w^1118,^*N=3, n=136), and (f’) *Hml^Δ^>GFP/UAS-bmm* (N=3, n=85, p=0.0004) in comparison to HSD *Control*, (*Hml^Δ^>GFP/w^1118,^* N=3, n=51). (e’’-f’’) Representative images of adult females on Ct.HSD from respective genetic backgrounds compared to control (d’).

**Figure S10:**
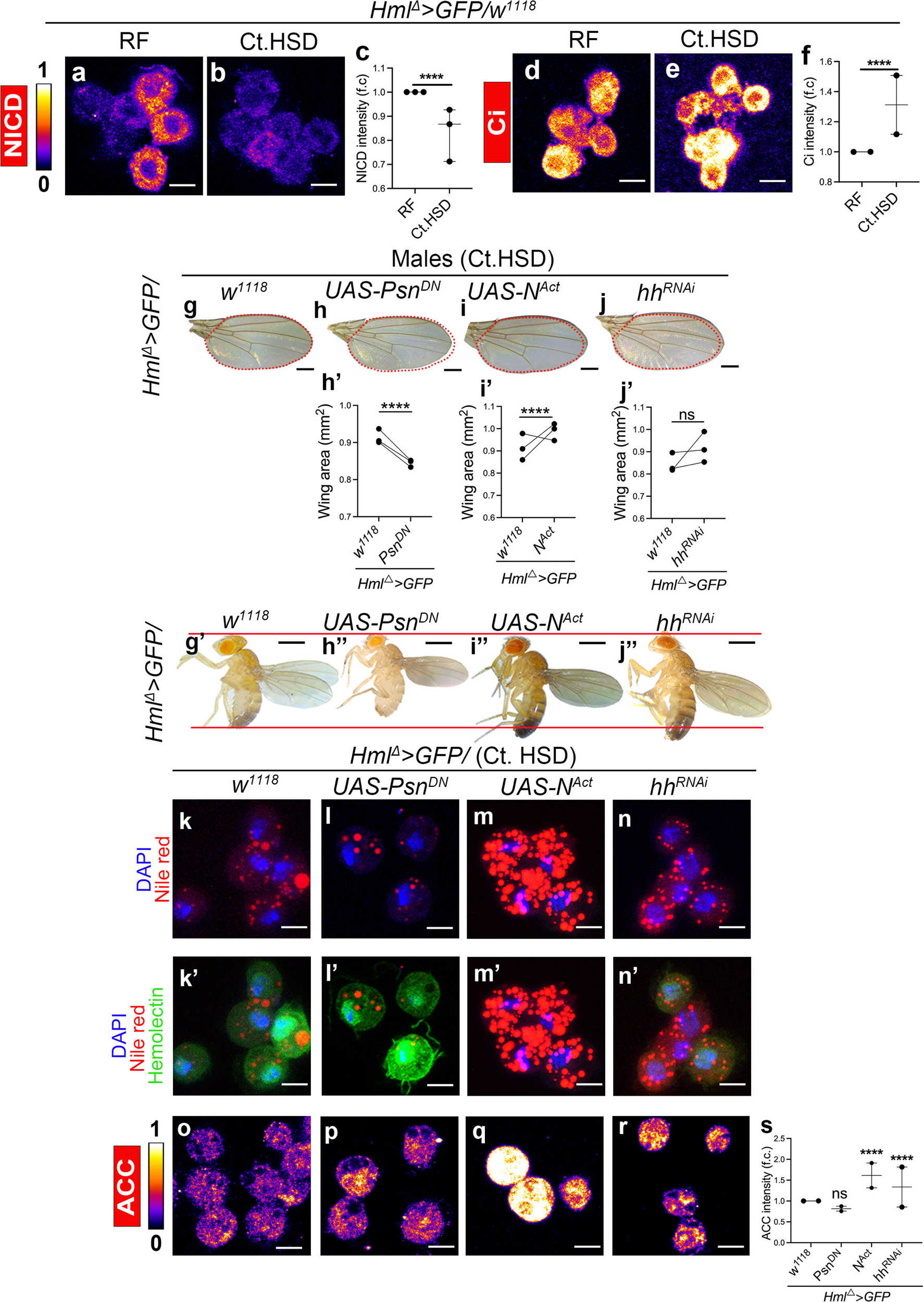
Hedgehog and Notch as upstream modifiers of immune cell lipogenesis in high sugar diet. Data information: DNA is stained with DAPI (blue), immune cells are shown in green (*Hml^Δ^>UAS-GFP*). NICD, Ci and ACC staining are shown in spectral mode in panels (a-b), (d-e) and (o-r). Nile red staining to mark lipids is shown in red in panels (k-n’). Scale bar: 5μm for immune cells, 0.5mm for flies and 0.25mm for wings. In quantification graphs (c), (f), (h’-j’) and (s), each dot represents an experimental repeat. Except for panels (a-b) and (d-e), where comparisons are with respect to Control on RF, in all other panels comparison for significance is with respect to Control on HSD. Asterisks mark statistically significant differences (*p<0.05; **p<0.01; ***p<0.001; ****p<0.0001). The statistical analysis applied is Two-way ANOVA, main effects or with Dunnett’s multiple comparison test wherever applicable. “N” is the total number of repeats. “n” is the total number of animals (larvae or adult flies) analyzed. Only right wing from each adult fly was selected for quantification. The red dotted line in the panels marks the wing span area and is used as a reference to showcase any change in wing span area across genotypes. The two horizontal red lines in the panels is used as a reference to showcase any change in fly lengths across genotypes. RF and Ct.HSD correspond to regular food and constitutive high sugar diet respectively. See methods for further details on larval numbers and sample analysis for each of the experiments. (a-b) Representative images of immune cells stained to visualize Notch Intracellular domain (NICD) expression. Compared to NICD protein levels in (a) RF Control (*Hml^Δ^>GFP/w^1118^*), (b) long-term HSD condition (Ct.HSD) showed a significant decrease. (c) Quantification of NICD intensity in RF (N=3, n=30) and Ct.HSD (N=3, n=30, p<0.0001). (d-e) Representative image of immune cells for active form of Cubitus interruptus (Ci) expression in Control (*HmlΔ>GFP/w^1118^*) regular food (RF) (d). Constitutive high sugar diet (Ct.HSD) (e) shows more Ci expression as compared to Control (d). (f) Quantification of Ci intensity in RF (N=2, n=30) and Ct.HSD (N=2, n=30, p<0.0001). (g-j’’) Modulating larval immune cell Notch and Hh pathway affects adult growth. Representative images of fly wings of adult males (g-j) showing size phenotype on Ct.HSD from respective genetic backgrounds. Compared to (g) Ct.HSD *Control* (*Hml^Δ^>GFP/w^1118^*), (h) loss of *Psn* (*Hml^Δ^>GFP/Psn^DN^*) or (i) increase in Notch activity (*Hml^Δ^>GFP/UAS-Notch^Act^*) in immune cells causes either a further reduction in adult size or recovery respectively. (j) Down regulating *hh* (*Hml^Δ^>GFP/hh^RNAi^*) in immune cells shows increased adult size compared to control (g). (h’-j’) Quantification of male wingspan in (h’) *Hml^Δ^>GFP/Psn^DN^*(N=3, n=73, p<0.0001) compared to Ct.HSD *Control (Hml^Δ^>GFP/w^1118,^*N=3, n=99,) (i’) *Hml^Δ^>GFP/UAS-Notch^Act^* (N=3, n=94, p<0.0001) compared to Ct.HSD *Control* (*Hml^Δ^>GFP/w^1118,^* N=3, n=119, p<0.0001), (j’) *Hml^Δ^>GFP/UAS-hh^RNAi^* (N=3, n=159, p=0.1313) compared to Ct.HSD *Control* (*Hml^Δ^>GFP/w^1118^,* N=3, n=72,). (h’’-j’’) Representative images of adult males on Ct.HSD from respective genetic backgrounds compared to control (g’). (k-n) Representative images of immune cells on Ct.HSD for lipids (Nile red) in (k) Ct.HSD *Control* (*HmlΔ>GFP/w^1118^*). (l) *UAS-Psn^DN^* (*HmlΔ>GFP/UAS-Psn^DN^*) and (m*) UAS-Notch^Act^* (*HmlΔ>GFP/UAS-Notch^Act^*) show decreased and increased lipids respectively as compared to Control (k). *hh^RNAi^* (*HmlΔ>GFP/hh^RNAi^*) (n) shows increased lipid droplets as compared to Control (k). (o-r) Representative images of immune cells for Acetyl CoA carboxylase (ACC) enzyme expression on (o) Ct.HSD *Control* (*HmlΔ>GFP/w^1118^*), *Psn^DN^* (*HmlΔ>GFP/UAS-Psn^DN^*) (p) shows no change as compared to Control. *UAS-Notch^Act^*(*HmlΔ>GFP/UAS-Notch^Act^*) (q) and *hh^RNAi^* (*HmlΔ>GFP/hh^RNAi^*) (r) show increased ACC expression as compared to Control (o). (s) Quantification of ACC intensity in *HmlΔ>GFP/w^1118^* (*Control*, Ct.HSD, N=2, n=30), *Psn^DN^* (*HmlΔ>GFP/UAS-Psn^DN,^*N=2, n=30, p=0.0945), *UAS-Notch^Act^* (*HmlΔ>GFP/UAS-Notch^Act,^* N=2, n=30, p<0.0001), *hh^RNAi^* (*HmlΔ>GFP/ hh^RNAi,^* N=2, n=30, p<0.0001).

**Figure S11.**
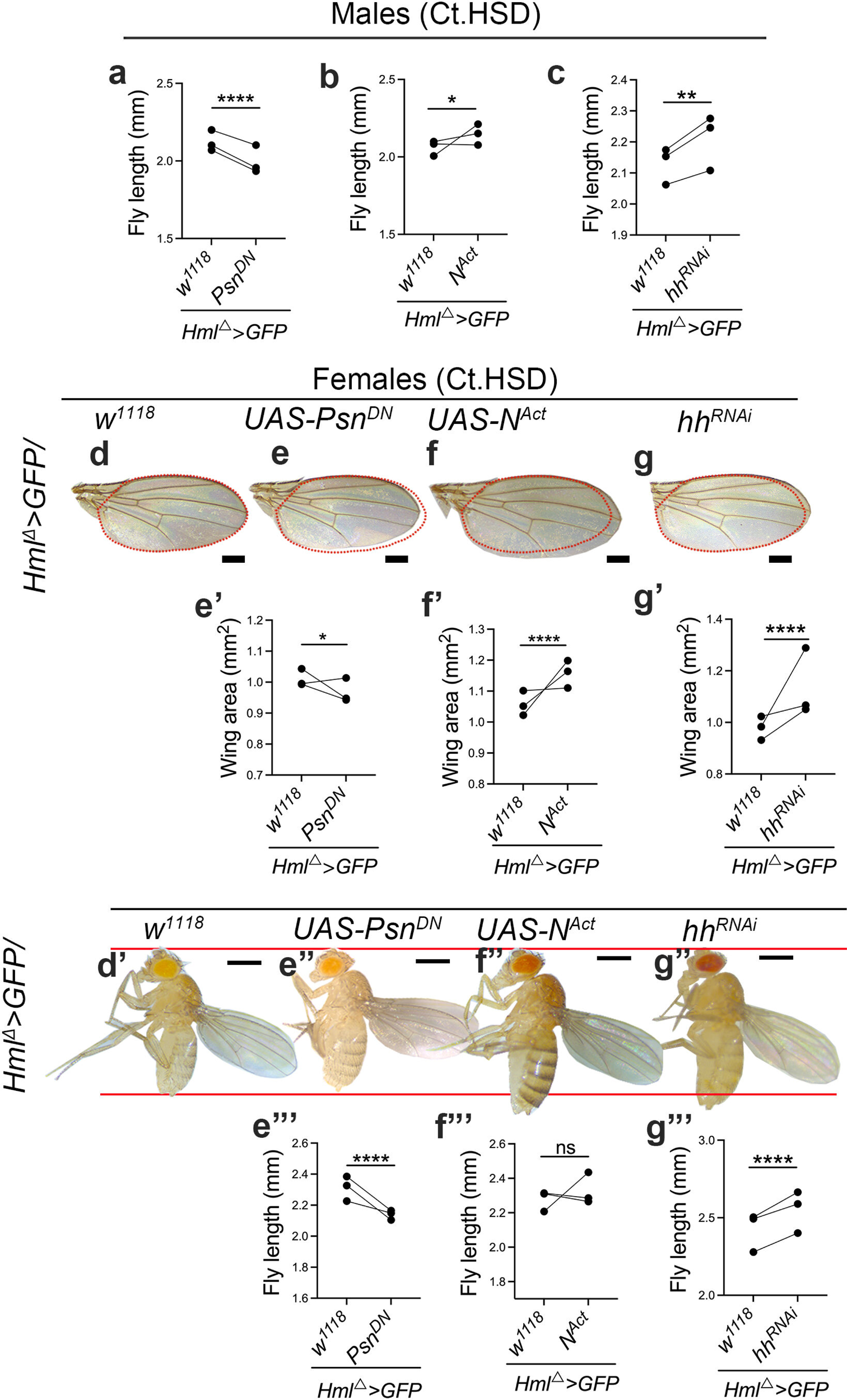
Hedgehog and Notch as upstream modifiers of immune cell lipogenesis in high sugar diet. Data information: Scale bar: 0.5mm for flies and 0.25mm for wings. In quantification graphs, shown in panel (a-c), (e’-g’), (e’’’-g’’’), each dot represents an experimental repeat. Comparison for significance is with respect to Control on HSD. Asterisks mark statistically significant differences (*p<0.05; **p<0.01; ***p<0.001; ****p<0.0001). The statistical analysis applied is Two-way ANOVA, main effects test. “N” is the total number of repeats. “n” is the total number of animals analyzed. Only right wing from each adult fly was selected for quantification. The red dotted line in the panels marks the wing span area and is used as a reference to showcase any change in wing span area across genotypes. The two horizontal red lines in the panels is used as a reference to showcase any change in fly lengths across genotypes. RF and Ct.HSD correspond to regular food and constitutive high sugar diet respectively. (a-g’’’) Modulating larval immune cell Notch and Hh pathway affects adult growth. (a-c) Fly length quantifications in males in (a) *Hml^Δ^>GFP/Psn^DN^*(N=3, n=51, p<0.0001) compared to Ct.HSD *Control*, (*Hml^Δ^>GFP/w^1118,^* N=3, n=58), (b) *Hml^Δ^>GFP/UAS-Notch^Act^* (N=3, n=40, p=0.0204) compared to Ct.HSD *Control*, (*Hml^Δ^>GFP/w^1118^* N=3, n=66,), (c) *HmlΔ>GFP/ Hh^RNAi^* (N=3, n=47, p=0.0012) compared to Ct.HSD *Control*, (*Hml^Δ^>GFP/w^1118^*N=3, n=58). Representative images of fly wings of adult females (d-g) showing size phenotype on Ct.HSD from respective genetic backgrounds. Compared to (d) Ct.HSD *Control* (*Hml^Δ^>GFP/w^1118^*), (e) loss of *Psn* (*Hml^Δ^>GFP/Psn^DN^*) or (f) increase in Notch activity (*Hml^Δ^>GFP/UAS-Notch^Act^*) in immune cells causes either a further reduction in adult size or recovery respectively. (g) Down regulating *hh* (*Hml^Δ^>GFP/hh^RNAi^*) in immune cells shows increased adult size compared to control (d). (e’-g’) Quantification of female wingspan in (e’) *Hml^Δ^>GFP/Psn^DN^* (N=3, n=58, p=0.0166) compared to Ct.HSD *Control*, (*Hml^Δ^>GFP/w^1118,^* N=3, n=60,) (f’) *Hml^Δ^>GFP/UAS-Notch^Act^* (N=3, n=72, p<0.0001) compared to Ct.HSD *Control*, (*Hml^Δ^>GFP/w^1118^* N=3, n=101,), (g’) *Hml^Δ^>GFP/hh^RNAi^* (N=3, n=141, p<0.0001) compared to Ct.HSD *Control*, (*Hml^Δ^>GFP/w^1118^*N=3, n=84). (e’’-g’’) Representative images of adult females on Ct.HSD from respective genetic backgrounds compared to control (d’). (e’’’-g’’’) Female body length quantifications in (e’’’) *Hml^Δ^>GFP/Psn^DN^*(N=3, n=46, p<0.0001) compared to Ct.HSD *Control*, (*Hml^Δ^>GFP/w^1118,^*N=3, n=50), (f’’’) *Hml^Δ^>GFP/UAS-Notch^Act^* (N=3, n=46, p=0.6726) compared to Ct.HSD *Control*, (*Hml^Δ^>GFP/w^1118^* N=3, n=56,), (g’’’) *HmlΔ>GFP/Hh^RNAi^* (N=3, n=28, p<0.0001) compared to Ct.HSD *Control*, (*Hml^Δ^>GFP/w^1118^* N=3, n=53).

